# Sensitivity to Vγ9Vδ2TCR T cells is imprinted after single mutations during early oncogenesis

**DOI:** 10.1101/2024.11.19.624272

**Authors:** Astrid Cleven, Angelo D. Meringa, Peter Brazda, Domenico Fasci, Thijs Koorman, Tineke Aarts, Inez Johanna, Dennis X Beringer, Patricia Hernandez-Lopez, Sabine Heijhuurs, Tomohiro Mizutani, Sangho Lim, Maarten Huismans, Jochem Bernink, David Vargas Diaz, Wei Wu, Esther San Jose, Jelle Schipper, Nikos Tsakirakis, Lauren Hoorens van Heyningen, Annick Nouwens, Lucrezia Gatti, Trudy Straetemans, Hugo Snippert, Jeanine Roodhart, Patrick W.B. Derksen, Jarno Drost, Maarten Altelaar, Albert J.R. Heck, Hans Clevers, Juergen Kuball, Zsolt Sebestyen

## Abstract

Vγ9Vδ2T cells have the unique ability to recognize a broad range of malignant transformed cells. The tumor targeting event involving BTN2A1 and BTN3A1 dimers on the tumor cell surface is critical, leading to full activation of the TCR. Although the molecular mechanisms governing TCR engagement and T cell activation are well-characterized, the role of Vγ9Vδ2 T cells in cancer immune surveillance remains to be fully elucidated, particularly the mechanisms that enable these cells to discriminate between healthy and malignant cells at an early stage of malignant transformation. We employed two independent, genetically engineered step-wise mutagenesis models of human colorectal and breast cancer that mimic the transformation steps leading to tumor formation. We demonstrate that various single oncogenic mutations introduced into healthy organoids or cells, are sufficient to upregulate surface expressed BTN2A1 and enable Vγ9Vδ2 TCR binding to tumor cells. However, full activation of T cells through a Vγ9Vδ2TCR required additional subsequent phosphorylation of juxtamembrane (JTM) amino acids of BTN3A1, leading to the activating heterodimerization of BTN2A1 and 3A1. Using a protein interactome mapping pipeline, we identified PHLDB2, SYNJ2 and CARMIL1 as key players in controlling these delicate dual surface dynamics of BTN2A1 and 3A1 during early transformation. This mode of action allowed Vγ9Vδ2TCR T cells to control tumors *in vitro* and in vivo, emphasizing the crucial role of these molecules from early mutagenesis, to advanced cancer stages, and highlighting the therapeutic potential of a Vγ9Vδ2TCR.

## INTRODUCTION

Increasing evidence shows that human γδT cells have an essential role in cellular stress sensing and immune surveillance, for both microbial and autologous stress (e.g. tumorigenesis)^1^. Infiltration of γδT cells in various tumors has been shown to have a favorable prognostic value^2^ and plays an important role in the immunosurveillance of early tumor development in mice^3^, indicating that γδT cells are most likely at the first line of defense during transformational processes of a healthy to a cancer cell. However, it remains unclear which process triggers γδT cells during early transformation, even though the anti-tumor role of γδT cells has been implicated in various tumor models with established tumors^4–7^. Vγ9Vδ2T cells, which are considered the most innate-like subset of gamma delta T cells in general^1^, are activated by intermediate metabolites of the isoprenoid/mevalonate pathway, such as isopentenyl pyrophosphate (IPP)^8^, also referred to as phosphoantigens (pAgs), which can build up in cancerous- or virally-infected cells due to disruption of the mevalonate pathway. Aminobiphosphonate (ABP) drugs, such as pamidronate (PAM), can also further increase cellular pAg levels, by inhibiting farnesyl diphosphate synthase (FPPS), an essential enzyme in this pathway^9^. Because of the broad, but tumor-specific recognition of malignantly transformed cells, and the lack of MHC-restriction, Vγ9Vδ2T cells harbor great clinical potential as an immunotherapy for cancer^10–12^. Although it is well-established that the γδTCR itself is essential for recognition of pAgs, the exact mechanism of ligand-receptor interaction has not been discovered, yet intracellular pAgs are bound to the B30.2 domain of butyrophilin-3 isoform A1 (BTN3A1) which leads to complex formation with BTN2A1^11,13–15^. BTN2A1 has emerged as a key protein for recognition of tumor cells by Vγ9Vδ2-T cells^14^ where it is directly bound by the gamma chain of Vγ9Vδ2TCR. The role of the small GTPase RHOB was previously demonstrated to relocalize to the membrane, and interact with BTN3A1 upon pAg build-up^16^. This is associated with cytoskeletal changes and reduced mobility of BTN3A1 in the membrane, implying a sort of “membrane trapping” mechanism. It is thought that direct binding of pAgs to the intracellular B30.2 domain enables heterodimerization of BTN3A1 with BTN2A1. The subsequently induced joint conformational and spatial changes stabilize BTN2A1-homodimers, which are then available for interaction with the γδ-TCR^16–18^.

Even with the identification of BTN2A1^11,13,14^, the understanding of BTN3A1 regulation through RHOB^11,16^, and the most recent observation that pAgs glue BTN3A1 and BTN2A1^18^, it is not fully elucidated how this mechanism is regulated. Recently, it has been shown to be partially controlled by the AMP-activated protein kinase (AMPK) pathway, and activation of this pathway led to increased transcription of both molecules, BTN3A and BTN2A1, followed by enhanced activation of Vγ9Vδ2T cells^19^. Although these new insights defined a signature which is predictive for recognition of tumors in cancer patients^19^, the signature could not clarify at which stage of transformation from and a healthy cell to a cancer cell the BTN-pathway is turned on. Therefore, we used a step-wise mutagenesis model for colon and breast cancer, which remodels different steps during mutagenesis, to characterize expression patterns of BTN2A1, BTN3A1 and RHOB during transformation, which identified PI3K activity as essential step to upregulate BTN2A1. This model also allowed us to hunt for novel players regulating, in particular, the heavily orchestrated BTN3A1 molecule by an innovative proximity proteomics approach. To properly analyze recognition of a tumor cell through Vγ9Vδ2TCR and overcome diversity in innate receptor expression and diversity in function of natural Vγ9Vδ2T cells, we used soluble Vγ9Vδ2TCR formats^11,12^ as well as αβT cells expressing a high affinity Vγ9Vδ2 TCR^20–22^. This strategy enabled us to characterize the orchestration of BTN2A1 and BTN3A1 during early transformation and late-stage cancers where BTN2A1 traffics early during mutagenesis to the cell membrane, in close proximity to BTN3A1, a process heavily regulated by PHLDB2, SYNJ2 and CARMIL1.

## RESULTS

### Vγ9Vδ2TCR T cells target early transformation events in colorectal cancer (CRC) and these features are preserved in late CRC stages to enable targeting by Vγ9Vδ2TCR T cells

We investigated both the *a priori* sensitivity of tumors to recognition by a Vγ9Vδ2TCR, as well as the presence of an altered mevalonate pathway in healthy versus diseased tissues by adding Pamidronate (PAM). In screenings to assess tumor recognition in the absence and presence of PAM, αβT cells expressing Vγ9Vδ2TCR exhibited the production of IFNγ when co-cultured with a variety of colorectal cancer (CRC) cell lines, and patient derived tumor organoids selectively in the presence of PAM **(Figure 1A)**. To discern whether recognition by a Vγ9Vδ2TCR is also a general mechanism of healthy colon tissues or a distinct feature of malignant transformation, we utilized a colon organoid model derived from normal tissues, which were altered to carry *APC* (APC^KO^), *KRAS* (KRAS^G12D^), *TP53* (p53^KO^) and *SMAD4* (SMAD4^KO^), referred to as AKPS mutant, simulating a fully developed CRC ^23^. IFNγ production after co-incubation of Vγ9Vδ2TCR T cells was exclusively observed in AKPS mutants in the presence of PAM, implying that recognition of tumors by a Vγ9Vδ2TCR is a hallmark of malignant transformation (**Figure 1B** and Supplementary Figure 1A-B). Vγ9Vδ2TCR T cells were also able to control CRC outgrowth for at least 43 days after tumor injection **(Figure 1C)** *in vivo,* and improved overall survival (**Supplementary Figure 1C**) when NSG-mice were engrafted with APKS mutant CRC organoids ^23^.

**Figure 1.**
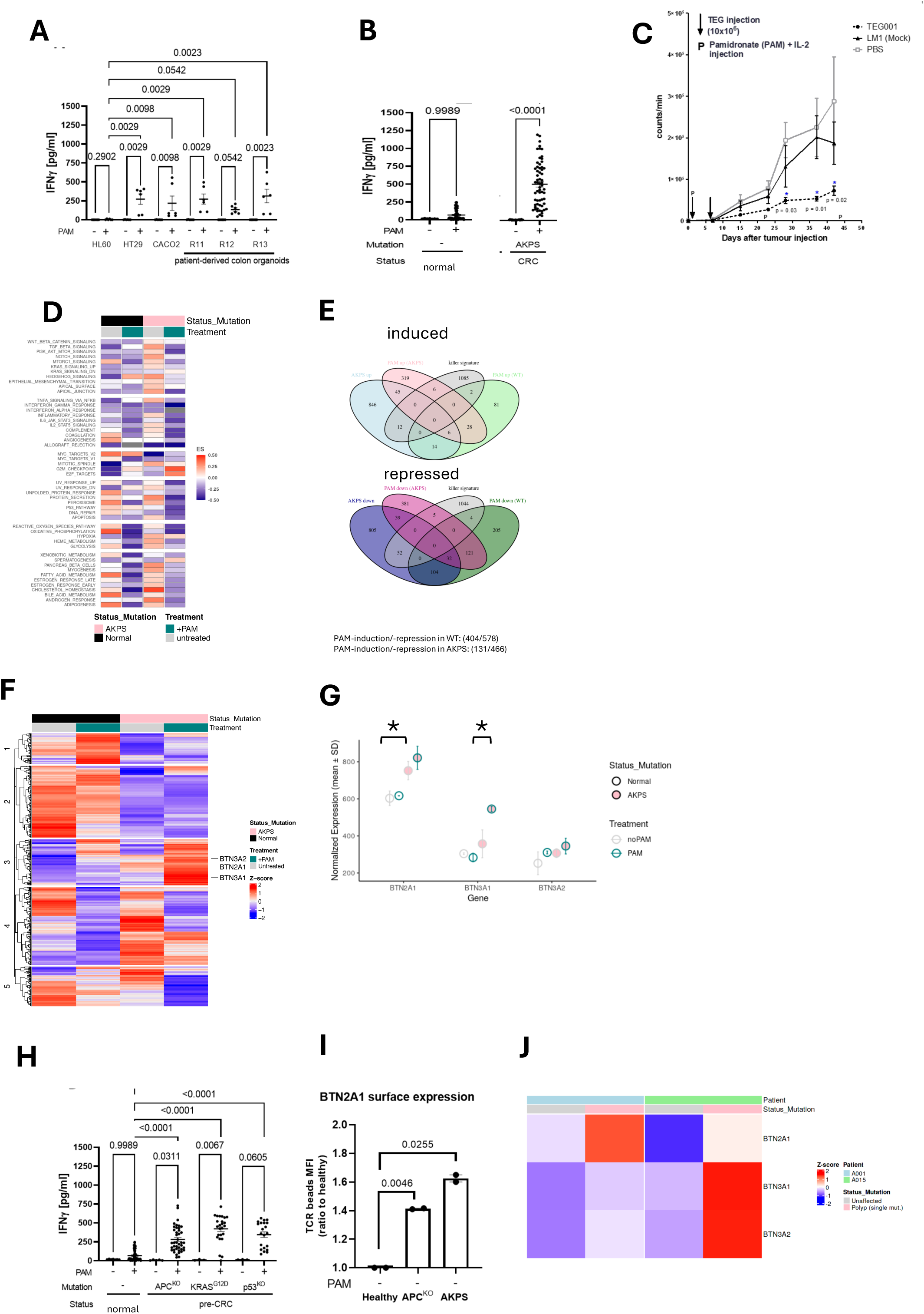
Vγ9Vδ2 TCR T cells recognize early transformed CRC organoids. **(A)** Patient derived CRC organoids (PDO) were co-cultured with Vγ9Vδ2TCR T cells in the presence of PAM and IFNγ release of T cells was determined by ELISA. **(B)** IFNγ production by Vγ9Vδ2TCR T cells after co-culture with either healthy colon organoids (normal) or CRC organoids mutated for APC, p53, KRAS and SMAD (AKPS) in the presence of 100uM PAM. **(C)** *In vivo* efficacy of Vγ9Vδ2TCR T cells against AKPS CRC organoids in the presence of PAM. Mice were treated with either PBS, T cells expressing a non-functional Vγ9Vδ2TCR (LM1) or a high affinity Vγ9Vδ2TCR (TEG001). Tumor burden of AKPS CRC organoids assessed by *in vivo* bioluminescence imaging (BLI) measuring integrated density per entire tumor area of mice. Statistical significances were calculated by mixed-effects model with repeated measures; *, P < 0.05; ***, P < 0.001. **<u>(I</u>)** IFNγ production by Vγ9Vδ2TCR T cells after co-culture with either healthy colon organoids (normal) or CRC organoids single-mutated for APC, p53 and KRAS in the presence of 100uM PAM. **(D)** Cluster heatmap displaying Gene Set Variation Analysis (GSVA) enrichment scores (ES) for HALLMARK gene sets in normal and AKPS samples, with or without 100µM PAM **(E)** Venn diagram illustrating differentially expressed genes (DEGs) with log fold change >1 and adjusted p-value ≤0.001, derived from pairwise comparisons of normal and AKPS samples, with or without 100µM PAM. **(F)** Heatmap representation of the expression of genes from the Murad ‘killer signature’ across wild-type and PAM +/- AKPS models. **(G)** Expression of BTNx genes in the WT-AKPS model, with and without PAM (normalized mean expression counts) **(H)** IFNγ production by Vγ9Vδ2TCR T cells after co-culture with either healthy colon organoids (normal) or CRC organoids single mutated for APC, p53 and KRAS in the presence of 100uM PAM. **(I)** Healthy colon organoids (WT), CRC organoids mutated for APC^KO^ or AKPS mutant CRC were stained with microbeads coated with either non-functional LM1 soluble Vγ9Vδ2TCR (LM1) or with high affinity soluble Vγ9Vδ2TCR (Cl5) in the absence of PAM. Data show MFI of bead binding**. (J)** Expression of BTNx genes in epithelial cells from adenocarcinoma patients, represented as per sample pseudobulk profiles from a publicly available single-cell RNAseq dataset.

To delineate whether sensitivity to Vγ9Vδ2TCR after PAM-treatment is already imprinted in organoids after malignant transformation, and whether PAM treatment acts differently on healthy versus malignant tissues, we conducted RNA-sequencing from the normal (healthy) and AKPS organoids, with and without PAM. We applied a Wilcoxon rank sum test over the enrichment scores of a gene set variation analysis (GSVA) focusing on HALLMARK gene sets **(Figure 1D)**. The test comparing normal and AKPS organoids without PAM treatment resulted in a p-value of 0.02312, indicating a statistically significant difference in pathway enrichment between WT and AKPS under ‘no PAM’ conditions, and implying that malignant transformation does create a different molecular imprint on cells. The major changes associated with the AKPS mutation covered biological terms related to signaling. We noticed that the highest enrichment score was calculated for the ‘CHOLESTEROL_HOMEOSTASIS’ term in the mutated samples, representing genes of the mevalonate pathway (ACAT, HMGCS1, MVK, PMVK, MVD, IDI1, FDPS).Comparing either healthy or AKPS-mutants with, or without PAM showed significant differences in enrichment scores in the normal (pval=0.0005891) and in the AKPS conditions (pval= 7.761e-09), mainly associated with the cell cycle, and implied that PAM acts on gene expression of both healthy and malignant tissues. However, when comparing the PAM-treated normal and PAM-treated AKPS-mutant organoids, there was no significant change in enrichment scores (p-value of 0.8822), suggesting that PAM’s effect on the gene expression landscape was consistent across both normal and AKPS conditions, as far as the HALLMARK scores are concerned, and that the different underlying molecular profiles in healthy and transformed tissues are the primary reason for PAM-responsiveness, and are not a selective effect of PAM on tumor cells only. Within this context, differential expression analysis comparing the effects of PAM treatment on gene activity demonstrated that normal organoids had a larger array of PAM-responsive genes, both induced and repressed (404 and 578 genes, respectively), than AKPS mutant organoids, which showed changes in 131 and 466 genes respectively **(Figure 1E)**. BTN genes, which are pivotal in the recognition of target cells by ψ8T cells^13,14^, demonstrated the most pronounced expression in AKPS mutants **(Figure 1G).** The BTN2A1 gene encoding for the main interacting protein with the Vγ9 chain, but not BTN3A1/2 genes, was already significantly elevated after malignant transformation in the absence of PAM, and was not further increased by PAM **(Figure 1G,)**. The addition of PAM significantly increased BTN3A1 gene expression levels in AKPS-mutants **(Figure 1G and Supplemental 1D).** However, patterns of other gene expression had little overlap with the ‘killer signature,’ a set of genes previously found in an extensive CRISPR-screening in the context of immune-mediated tumor cell destruction upon PAM treatment ^19^, implying that the recently published killer signature does not fully explain the general sensitivity upon malignant transformation to Vγ9Vδ2TCR-mediated recognition.

To assess at which stage malignant transformation can be made susceptible to Vγ9Vδ2TCRs, we made use of a step-wise mutagenesis colon organoid model, where the introduction of single mutations of *APC* (APC^KO^), *KRAS* (KRAS^G12D^), *TP53* (p53^KO^) and *SMAD4* (SMAD4^KO^) into normal (healthy) colon organoids represented models for pre-cancerous lesions, while the AKPS mutant served as a fully developed CRC^23^ model. After co-incubation of Vγ9Vδ2TCR T cells with organoids treated with/without PAM, a single gene mutation introduced in the normal colon was already able to induce IFNγ production after addition of PAM (**Figure 1H and Supplementary Figure 1A-B)**. We chose APC^KO^ and AKPS mutant CRC organoids to represent very early and late stages of CRC development, respectively, to further characterize the capacity of a Vγ9Vδ2TCR to potentially sense early or late-stage cancer development. On both APC^KO^ and AKPS mutants, we found an enhanced expression of BTN2A1 protein at the cell membrane in the absence of PAM (**Figure 1I**). Also, the PAM-induced redistribution of intracellular RhoB which we previously described,^11,16^ was already detected in single *APC*-mutant organoids (**Supplementary Figure 1E).**

To validate our observation that early malignant transformation primarily impacts BTN2A1 expression in pre-cancerous lesions in humans, we consulted existing single cell RNA-sequencing (scRNAseq) datasets from patients at various disease progression stages of CRC^24^. Analysis of pseudo bulk profiles of the epithelial cell population revealed that BTN2A1 expression levels were always heightened in pre-cancerous (polyp) stages when compared to the “unaffected” baseline in both of the studied patients (**Figure 1J**), which parallels the increased BTN2A1 expression seen at comparable stages in our model. We could also observe that in one pre-cancerous lesion, BTN2A1, BTN3A1 and BTN3A2 were increased.

### Phosphoinositide 3-kinase/AKT1 activity during early transformation is a prerequisite of BTN2A1 upregulation

To confirm that single oncogenic mutations generally mediate susceptibility to Vγ9Vδ2TCR recognition, we tested the effect of single *ERBB2* mutations, and overexpression in breast tissue^25–27^. MCF10a, a non-transformed breast epithelial cell line, was engineered to mimic various oncogenic *ERBB2* gene related mutations such as overexpression of HER2 (ErbB2^AMP^), a single mutation in the extracellular domain (ErbB2^S310F^) or amplified kinase activity mutant (ErbB2^V777E^). Only MCF10a cells expressing kinase mutant HER2 variant (ErbB2^V777E^) triggered IFNγ production of Vγ9Vδ2TCR T cells, again exclusively in the presence of PAM (**Figure 2A**), while benign cells and the two other oncogenic variants remained unrecognized. This suggests that it is not the transformation event in general that promotes susceptibility to recognition by a Vγ9Vδ2TCR, but rather very specific mutations that also differ, depending on the cellular phenotype of the tumor. Since mutations in *ERBB2* often result in PI3K pathway activation ^28^ we pre-treated MFC10a ErbB2^V777E^ tumor cells with PI3K inhibitor (Pictilisib), which led to significant reduction of IFNγ production of Vγ9Vδ2TCR T cells upon co-culture, while HER2 CAR-T cell reactivity remained unaffected **(Supplementary Figure 2A**). Surface expression of the Vγ9Vδ2TCR ligand, BTN2A1, was significantly increased upon ErbB2^V777E^ mutation in the absence of PAM. This increase was dependent on PI3 kinase activity in MCF10a cells, since pan PI3K inhibition depleted BTN2A1 surface expression **(Figure 2B)**. BTN3A surface expression, however, did not change either upon ErbB2^V777E^ mutation or PI3K inhibition, but it significantly decreased upon any oncogenic mutations induced into benign cells (**Figure 2C**). We could confirm the dependence of BTN2A1 and independence of BTN3A1 expression, respectively, on PI3K signalling in multiple targeted cell lines in the absence of PAM **(Supplementary** Figures 2B and C). When analysing phosphorylation of AKT in MCF10a HER2 mutants, we found that while AKT was phosphorylated on position Ser473 in all lines, in the presence of full culture medium containing growth factors, only a mutation in position V777E had already induced phosphorylation in the absence of growth factors **(Figure 2D),** confirming the special role of this mutation. To further clarify the downstream signalling cascade of PI3K that is necessary for PAM-dependent activation of Vγ9Vδ2TCR T cells, we blocked various key kinases that might be activated by PI3K in the subsequent experiments. In all tumor cell lines, inhibition of both PI3K and AKT1 significantly reduced PAM-dependent Vγ9Vδ2TCR activity, while specific MEK inhibitors did not alter recognition of the tumor cells (**Figure 2E**). Pre-treatment of epithelial (HT-29, SKBR3) and hematological (Daudi, K562, RPMI-8226) tumor cell lines with blocking reagents of mTOR reduced recognition significantly, with a similar or even higher potency, as inhibiting AKT activity **(Figure 2F)**. Inhibition of both mTOR-involved complexes, mTORC1 and mTORC2, by inhibitor Torin1, was superior in reducing activation of Vγ9Vδ2TCR T cells compared to inhibition of only mTOR complex 1 (mTORC1) by Rapamycin **(Figure 2F)**. Since BTN2A1 surface expression was solely dependent on PI3K activity in tumor cell lines, while Vγ9Vδ2T cell activation was triggered by the combination of increased PI3K pathway activity and the presence of PAM, we wondered whether the activity of this pathway is exclusively transformation dependent. The expression patterns of genes of the PI3K-AKT-mTOR pathway over our AKPS-models displayed varied trends in terms of being induced by PAM, or by transformation. However, the key components of the pathway (PRKCA/B, PIK3CA, PAK1, AKT2) were predominantly influenced by the transformation process itself **(Figure 2G)** and thus highly correlated with BTN2A1 expression **(Figure 2H)**, suggesting that as the disease progresses, these key pathway components and BTN2A1 expression are linked. These combined data indicate that PI3K/AKT/mTOR activity in the early onset of malignant transformation drives BTN2A1 surface upregulation on tumor cells, and therefore makes them susceptible for Vγ9Vδ2TCR targeting.

**Figure 2.**
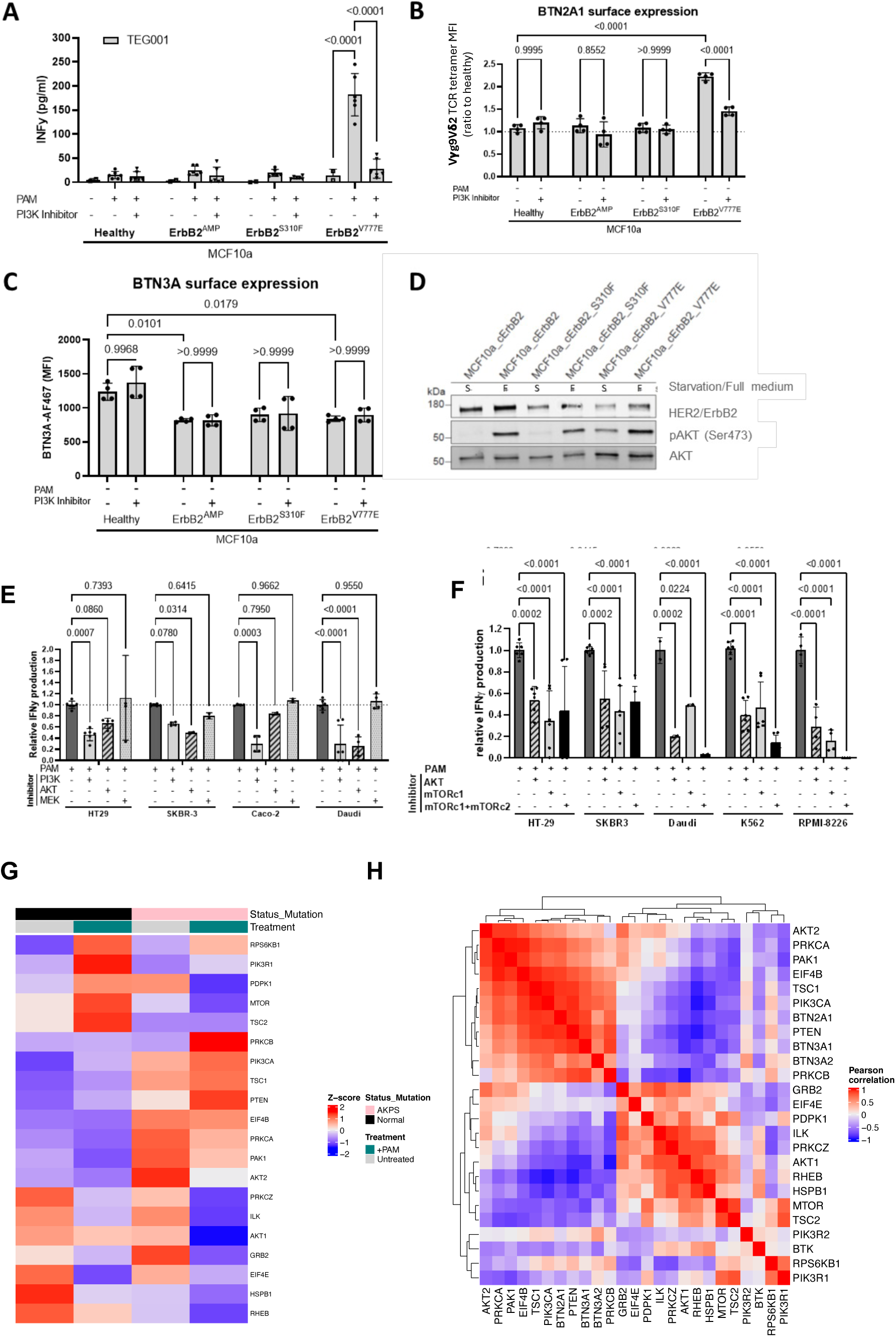
Transformed cells upregulate BTN2A1 surface expression via PI3K kinase activity. **(A)** MCF10a cells were transduced with different ErbB2 variants and co-cultured with either Vγ9Vδ2TCR or HER2-CAR transduced T cells. Tumor cells were pre-treated with the PI3K kinase inhibitor Pictilisib at 2uM overnight. After an overnight co-culture, supernatant was used to determine IFNγ production by the Vγ9Vδ2TCR T cells. **(B)** Furthermore, tumor cells were isolated and stained for BTN2A1 via TCR tetramer staining and **(C)** BTN3A cell surface expression. **(D)** Protein expression of HER2, phosphorylated AKT (pAKT) and total AKT in MCF10a mutant lines cultured for 24h in full culture medium *(F)* or medium without additional growth factors *(S)*. **(E)** Multiple tumor cell lines were co-cultured with Vγ9Vδ2TCR T-cells after pre-treatment with either the PI3K kinase inhibitor, AKT inhibitor or MEK inhibitor. After an overnight co-culture, supernatant was used to determine IFNγ production by the Vγ9Vδ2TCR T cells. **(F)** Multiple tumor cell lines were pre-treated with either PI3K kinase inhibitor, AKT inhibitor, mTOR inhibitor Rapamycin or mTOR inhibitor Torin1 and subsequently co-cultured with Vγ9Vδ2TCR T cells. After an overnight co-culture, supernatant was used to determine IFNγ production by the Vγ9Vδ2TCR T cells. **(G)** Heatmap illustrating the expression of genes from the ‘PI3K_and_AKT_family’ gene list, in the WT-AKPS model, with or without PAM. **(H)** Heatmap of Pearson’s correlation of the expression of genes from the ‘PI3K_and_AKT_family’ and BTNx genes in the WT-AKPS model, with or without PAM.

### BTN3A1 phosphorylation is required for full activation of the Vγ9Vδ2 TCR

To this end we revealed that transformation induced AKT/mTOR activity promotes cell surface expression of BTN2A1, which is a prerequisite for binding of the Vγ9 TCR chain. However, in all models, full activation of Vγ9Vδ2TCR T cells required PAM treatment. Although we have not been able to identify the driver of PAM-mediated BTN3A1 upregulated (Figure 1G) we could further characterize molecular mechanisms once BTN3A1 is upregulated, by revisiting previously generated mass spectrometry data generated to assess PAM-induced changes in surface proteins ^11^. This analysis identified two potential phospho-sites on the juxtamembrane (JTM) region of the BTN3A1 molecule; the residues S296 and T297 that were phosphorylated upon PAM **(Figure 3A**). The juxtamembrane region of BTN3A1 is important for Vγ9Vδ2 T cell activation, and is predicted to form a coiled-coil dimer, with the JTM of either BTN3A2 or BTN3A3. This heterodimerization is also important for the surface expression of BTN3A1 ^29^, but also plays a key role in the interaction of BTN3A1 with RhoB ^16^. In order to determine whether phosphorylation of S296 and/or T297 could contribute to interhelical electrostatic interactions, and thereby favor heterodimerization, a coiled-coil model of the 3A1 and 3A2 JTM was generated. Neither S296 nor T297 are located at the interface, and there are no positively charged residues located at the interface of the 3A2 helix (Fig 3B). As enhanced coiled-coil stability of the phosphorylated residues is less likely, the alternative of providing an improved scaffold for RhoB to bind, could be a valid alternative. To separate the functional impact of PAM-induced phosphorylation on BTN3A1 from other events induced by endogenous phosphoantigen (pAg) levels, we constructed modified versions of the BTN3A1 sequence that mimic protein conformation of either a phosphorylated (phospho-mimic variant) or an unphosphorylated state (phospho-deficient variant). These phospho-variants were developed by substituting residues S296 and T297 with aspartic acid, which is known to function similarly to phosphorylated serine and threonine (D, phospho-mimic), or alanine which is employed regularly to inhibit residue phosphorylation (A, phospho-deficient) To understand whether phosphorylation mimicking of the residues investigated here is required for BTN3A1 membrane orchestration, we performed FRET measurements that showed that phospho-deficient BTN3A1 is unable to interact with RhoB, another important player in the orchestration of BTN3A1^16,30^ **(Figure 3E)** suggesting a significant role of JTM phosphorylation of BTN3A1 in enabling Vγ9Vδ2TCR full activation., Functional introduction of the phospho-mimic BTN3A1 variant resulted in higher activation of Vγ9Vδ2TCR T cells upon PAM **(Figure 3C and 3D)**. To assess potential regulators of this phosphorylation we used gene expression profiles of the normal and AKPS-mutated colorectal organoids, and identified protein kinase C theta (PRCKQ) as the only kinase that follows PAM-treatment, and upregulates by PAM, regardless of the mutational status of the tissue (**Figure 3F**). This data led to the hypothesis that PAM-induced kinase activation results in phosphorylation of the S296 and T297 motifs of BTN3A1. BTN3A1 phosphorylation then alters interaction of BTN3A1 with RhoB.

**Figure 3.**
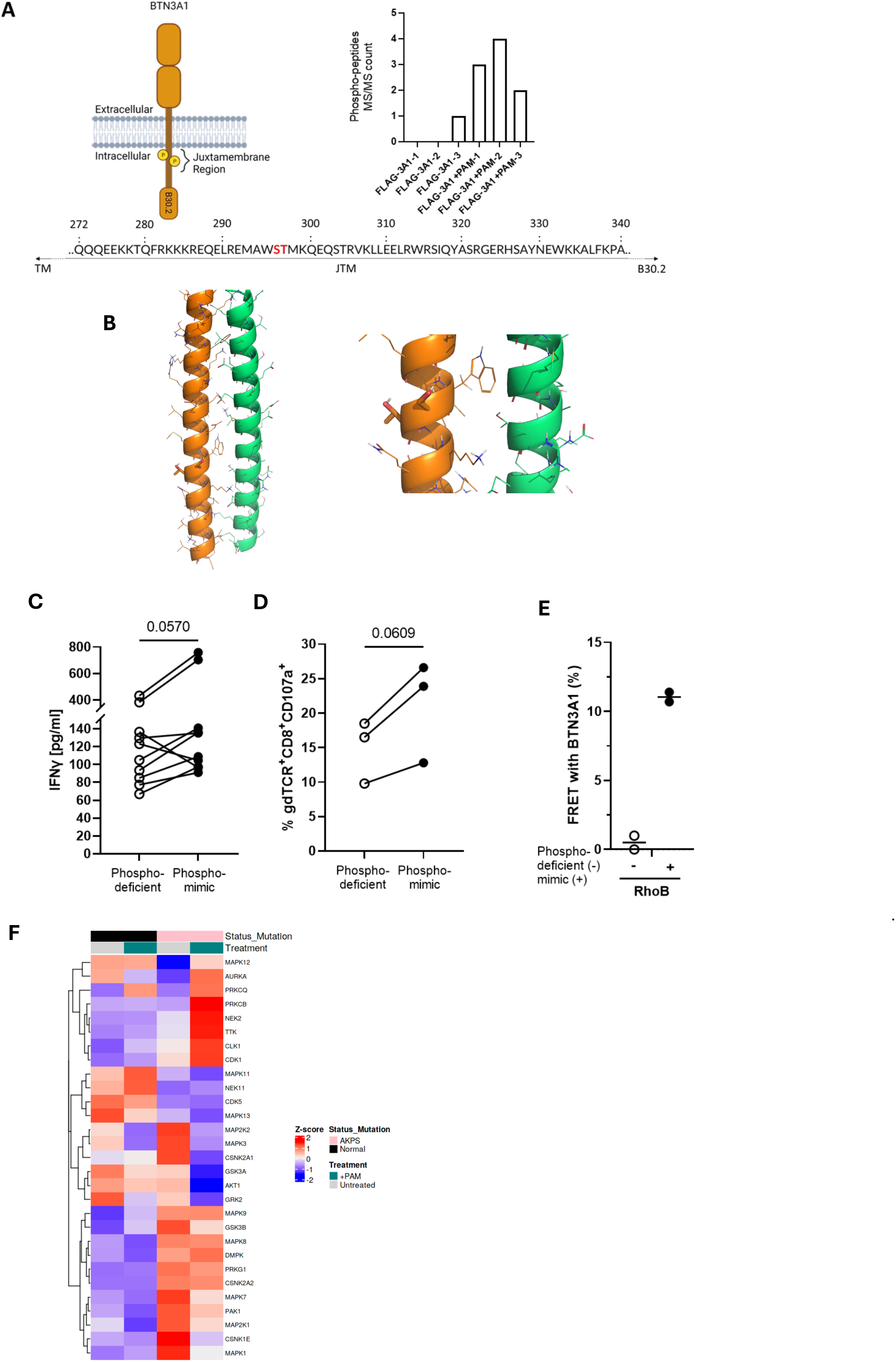
Phopshosites of juxtamembrane region of BTN3A1 affect Vγ9Vδ2TCR T cells target recognition. **(A)** Putative BTN3A1 phosphosites identified in previously published AP-MS proteomic data [REF. Vyborova et al.]. The bar plot indicates the number of times a peptide with a phosphorylation on S296/T297 or S326 was detected on FLAG-BTN3A1 immunoprecipitates from cells untreated or treated with PAM. **(B)** Predicted position of phosphor sites on BTN3A1-BTN2A1 heterodeimer. **(C)** IFNγ production of Vγ9Vδ2TCR T cells was measured upon co-culture with either the Phospho-deficient or the phospho-mimic mutated HEK293FT cells in the presence of PAM. **(D)** CD107a degranulation of Vγ9Vδ2TCR T cells was measured upon co-culture with either the Phospho-deficient or the phosphor-mimic mutated HEK293FT cells in the presence of PAM. **(E)** Flow cytometry FRET measurement was performed between phospho-mutated variants of BTN3A1 and RHOB. **(F)** Heatmap illustrating the expression of kinases predicted to be active targeting the S297 and T297 depicted by Piero Giansanti et.al., in the WT-AKPS model, with or without PAM.

### Pamidronate induces BTN3A1-proximate spatial protein interactome in tumor cells that is required for Vγ9Vδ2TCR-induced tumor targeting

To study in more detail which additional protein complexes are in the proximity of BTN3A1 after addition of PAM, we set up an interactome pipeline in the proximity of the BTN3A1 that are exclusively related to Vγ9Vδ2TCR activation (**Figure 4A**). As a first step, we made use of a proximity-dependent biotin identification (BioID) approach^31^ by overexpressing BTN3A1 fused to bacterial biotin ligase BirA* to its the C-terminus in HEK293F cells. Cells were treated with 100 µM PAM and 50 µM biotin for 24h, which led to biotinylation of BTN3A1-close proteins **(Supplementary Figure 4A** and B), after which cells were lysed, and biotinylated proteins were pulled down, using streptavidin beads. Proteins were identified by mass spectrometry, analyzing quadruplicate samples, which resulted in a list of 60 candidate proteins including 31 hits under no treatment, 19 upon PAM treatment, and 10 hits that occurred upon both conditions (**Supplementary Figure 4C)**. To determine if candidate proteins play a physiological role in tumor targeting by Vγ9Vδ2TCR T cells, we knocked down each candidate protein in HEK293F and MZ1851RC cells, using siRNAs and used these cells as targets against Vγ9Vδ2TCR T cells in the presence of PAM, and calculated to what level KD influenced IFNγ production by Vγ9Vδ2TCR T cells **(Figure 4B)**. We used the same target cells against Wilms tumor 1 specific (WT-1) αβTCR T cells after loading them with WT-1 peptide, to exclude that candidate proteins are not involved in general T cell-target cell interactions **(Supplementary Figure 5B)**. The summary of these knock-down experiments provided evidence that eight candidate proteins, namely PHLDB2, PKP2, SYNJ2, CARMIL1, PA2G4, TANC1, SDK2, HNRPC, have a functional impact on PAM-dependent activation of Vγ9Vδ2TCR T cells, as possible additional modulators of the BTN3A1-RhoB axis.

**Figure 4.**
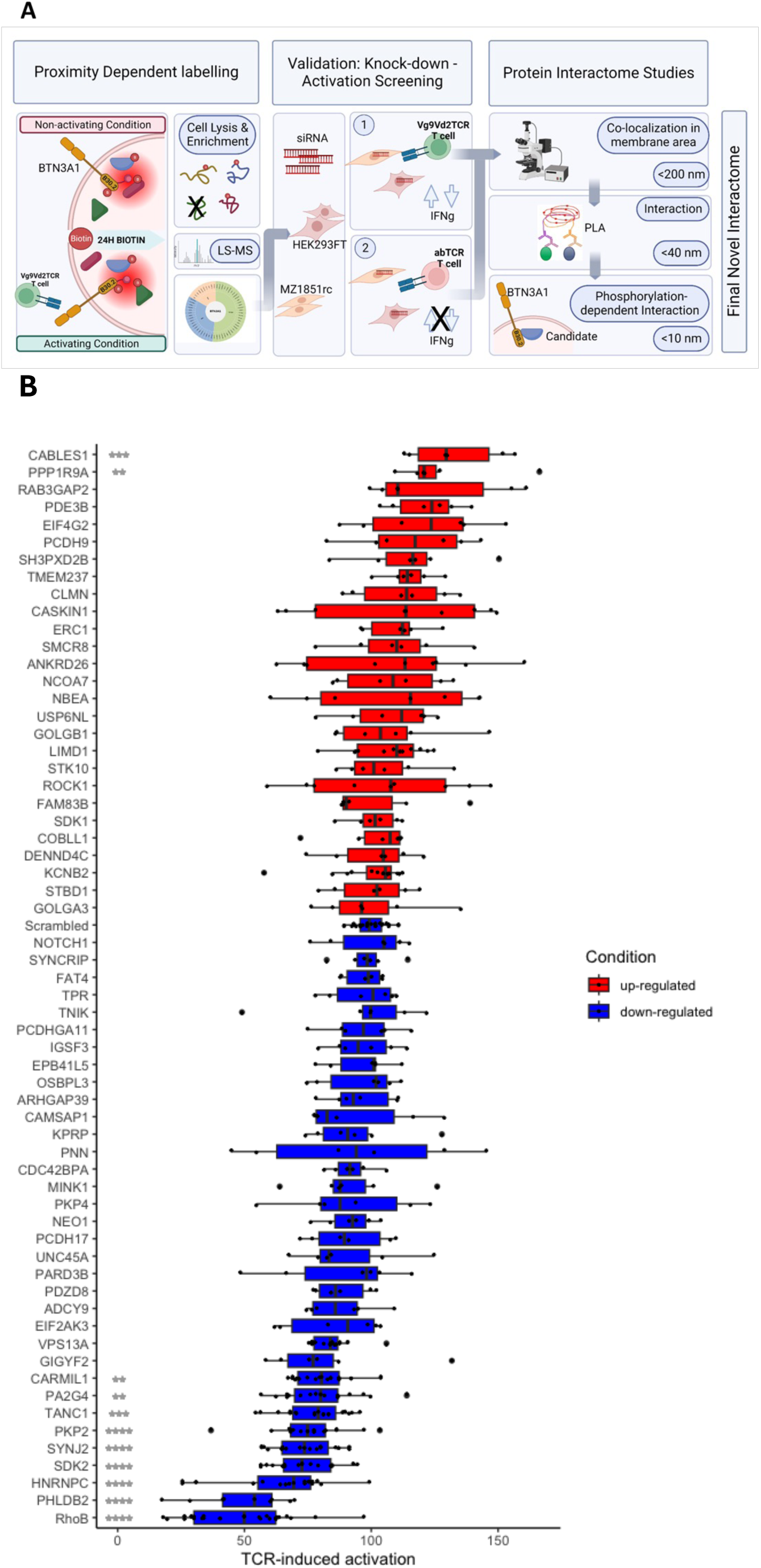
BioID identifies BTN3A-interacting proteins involved in T-cell tumor targeting. **(A)** Schematic representation of the BTN3A1 proteome characterization pipeline. **(B)** Genes as indicated have been transiently knocked-down using siRNA. After 48h Vγ9Vδ2TCR T cells and 100 µM PAM were added. After overnight co-culture, INFγ production of Vγ9Vδ2TCR T cells was measured. **(B)** Box-and-whisker plots depicting the levels of activation after siRNA knock-downs. Percentage of activation was normalized at each experiment to wild type target cells HEK 293T or MZ1851rc. Each gene is ranked according to the fold change in mean activation level relative to the control condition denoted as ‘Scrambled’. The genes are displayed in descending order of this fold change. Color coding indicates the direction of change compared to the ‘Scrambled’ baseline. (median is represented in box-and-whisker plots). A linear model (lm()) was employed to assess the differences in activation levels across the (knocked-down) genes. To dissect the differences between each gene and the ‘Scrambled’ control, estimated marginal means (EMMs) were calculated for each gene using the emmeans() function from the emmeans R package. Subsequently, pairwise comparisons were conducted employing Dunnett’s test. The results from these pairwise comparisons were summarized to include confidence intervals and adjusted p-values: Each comparison was annotated with significance markers to visually denote the level of statistical significance: **** for p < 0.001, *** for p < 0.01, ** for p < 0.05, * for p < 0.1, empty (ns) for non-significant results.

We wondered if PAM-induced spatial rearrangements of BTN3A1-related proteins, as observed with BioID-tagging, could be partially the result of post-transcriptional or post-translational regulation of proteins in tumor cells. Therefore, we assessed changes of expression of the candidate proteins upon PAM treatment in a panel of Vγ9Vδ2TCR-activating tumor cell lines, including breast cancer line MDA-MB 231, HEK293FT, renal cell carcinoma cell line MZ1851rc and head and neck cancer cell line SCC9, by performing Western blot analysis. In addition to BTN3 and RhoB we focused our further analysis on five out of eight new candidate proteins, against which reliable antibodies were available (PHLDB2, PKP2, SYNJ2, CARMIL1, PA2G4) that allowed both Western blot and cellular expression analysis. We found that PAM treatment induced a significant increase of RhoB and SYNJ2 protein, and decrease of PHLDB2 protein expression in all tested cell lines in the presence of PAM **(Figure 5A and Supplementary Figure 5D)**. Next, we compared protein expression in HEK293FT wt and HEK BTN3 KO cells in the presence, and absence of PAM, of the newly identified proteins, and observed a partial loss of expression of SYNJ2, suggesting that its expression is partially co-regulated with BTN3A1 **(Supplemental Figure 5E)**. To further map how BTN3A interaction dynamics occur with our new five candidate proteins in a PAM-dependent manner, we analyzed co-localization between BTN3A membrane clusters and each of the candidate proteins, using confocal microscopy in MZ1851RCcells. Examples of the region of interest selections and representative images are shown in (**Supplementary Figure 6A** and B). We found that while PHLDB2 moves out of BTN3A membrane clusters upon PAM treatment, SYNJ2 and CARMIL1 moves into them, within the range of approximately 200 nm **(Figure 5B).** To confirm if PHLDB2, SYNJ2 and CARMIL1 truly interact with BTN3A1 in tumor cell membranes, we studied interaction at higher resolution, measured by Proximity ligation assay (PLA) which can visualize protein interactions within approximately 40 nm. Similarly to the previously described small GTPase RHOB ^16^, SYNJ2 and CARMIL1 both showed a significant PAM-dependent interaction with BTN3A1, while PHLDB2 showed no significant interaction with BTN3A1 **(Figure 5C and Supplementary Figure 6C)**. To understand whether next to spatial regulation at the protein level, the expression of the genes investigated here is regulated similarly to BTNs, we examined the AKPS mutant CRC model gene expression data. While CARMIL1 and RHOB gene expression strongly correlated with BTN3A1 gene expression both in our normal colon and AKPS CRC models, PHLDB2 and SYNJ2 expression did not seem to be linked to these changes, suggesting that the involvement of these molecules in BTN3A1 expression is regulated entirely post-transcriptionally **(Figure 5D).** To demonstrate that the BTN3A1-network identified here is significant for the immunosurveillance of Vγ9Vδ2TCR T cells in a clinically more relevant setting, we knocked out the individual candidate genes in AKPS mutant organoids using the CRISPR/Cas system, and tested Vγ9Vδ2TCR T cells reactivity against them. KO of each individual gene similarly reduced both IFNγ production and direct tumor killing of Vγ9Vδ2TCR T cells in response to AKPS mutant organoids **(Figure 5E and F).**

**Figure 5.**
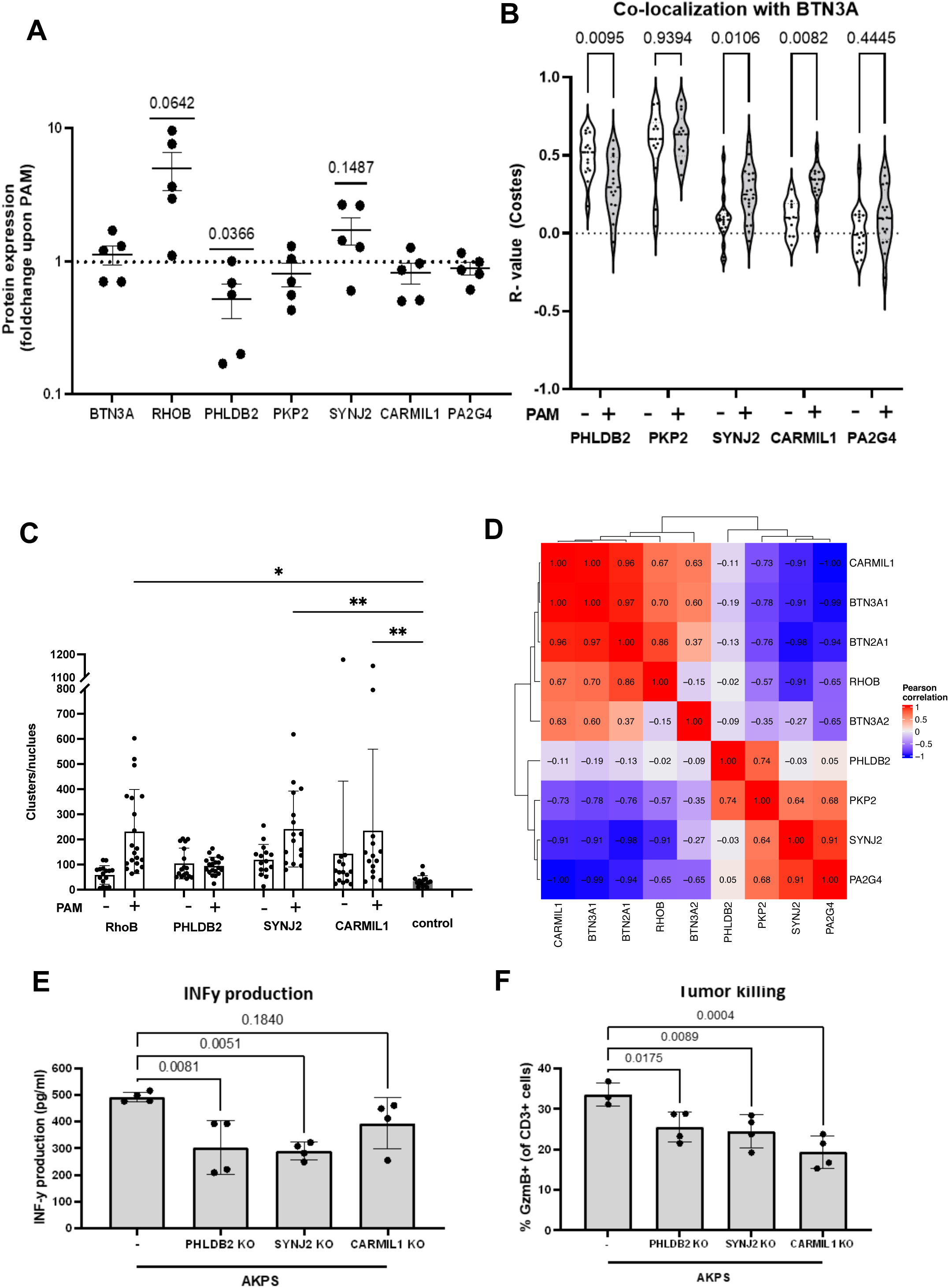
Candidate proteins differentially colocalize with BTN3A in membrane clusters. **(A)** Protein expression on 5 different targeted tumor cell lines was determined in the absence and presence of PAM by Western Blot, expression differences (PAM vs no PAM) as ratios are depicted in the figure**. (B)** MZ1851rc cells were treated overnight with either 0µM or 100µM PAM. Afterwards, cells were fixed, permeabilized, and stained for all of the indicated candidate interacting proteins and BTN3A1. Bars indicate mean +/-SEM. Statistical significance of differences between no PAM and PAM conditions were determined using unpaired parametric T-tests. All analyses were performed blinded to sample conditions. **(C)** MZ1851rc cells were either treated overnight with 100µM PAM or left untreated. Subsequently, cells were fixed and permeabilized. A DuoLink^TM^ proximity ligation assay (PLA) was performed to assess interaction between CD277 and either PHLDB2, SYNJ2, or CARMIL1 respectively. Each condition was paired with a technical control (C) constituted by leaving out one of the primary antibodies. All the technical control samples were pulled together to form the control condition. Multiplicity adjusted P-values were calculated using a two-way ANOVA with Tukey’s multiple comparison test. Bars indicate mean +/-SEM. **(D)** Heatmap of Pearson’s correlation of the expression of genes from the ‘BioID candidates’ and the BTNx genes in the WT-AKPS model, with or without PAM. **(E)** IFNγ production of Vγ9Vδ2TCR T cells after co-culture with AKPS CRC organoids knocked out for PHLDB2, SYNJ2 and CARMIL1, respectively in the presence of 100 µM PAM. **(F)** Granzyme B production of Vγ9Vδ2TCR T cells after co-culture with KO-variants of AKPS mutant organoids knocked out for PHLDB2, SYNJ2 and CARMIL1 with the presence of 100 µM PAM.

We conclude that PHLDB2, SYNJ2 and CARMIL1 are are involved in orchestrating PAM-induced cytoskeletal rearrangements in tumor cells that are a prerequisite for spatial and conformational changes in BTN3A1 leading to Vγ9Vδ2TCR T cell activation **(Figure 6).**

**Figure 6.**
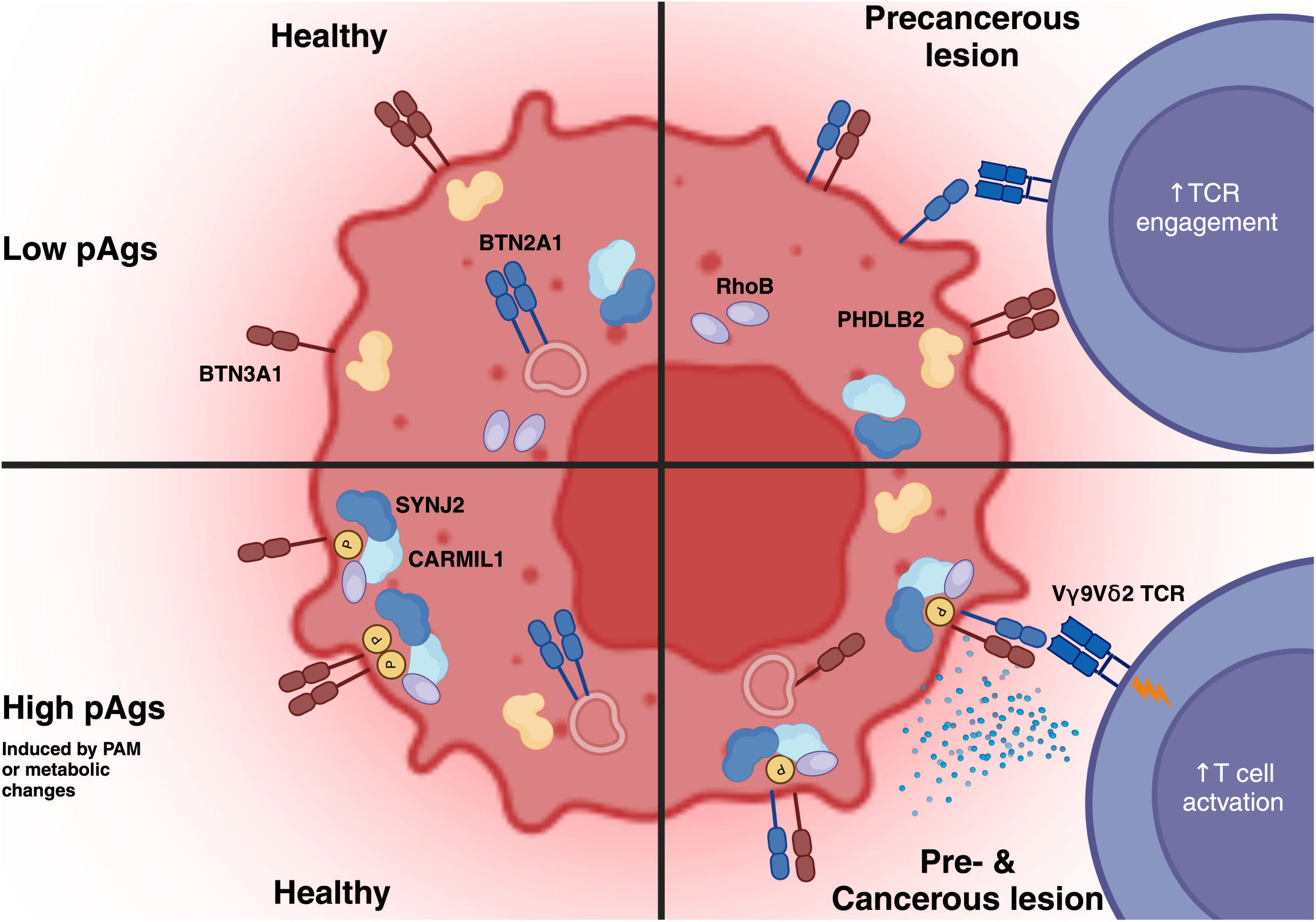
Schematic illustration of molecular steps needed for full activation of Vγ9Vδ2TCR T cells. The left panel represents healthy tissues and the right panel precancerous and cancerous tissues. The top panel shows molecular events at low intracellular phosphoantigen (pAgs) levels, while the lower panel represents high pAgs induced either by ABPs or by metabolic changes.

## DISCUSSION

Despite the growing interest in Vγ9Vδ2T cells and their anti-tumor potential, and the research aiming to resolve the unknown mechanisms of induction and regulation of this anti-tumor effect, there are still many mysteries to resolve. One of these is how tumor recognition is triggered, and another is how exactly it is regulated.

Here, we have demonstrated that potential sensitivity to the Vγ9Vδ2TCR T cells is imprinted as early as after single mutations in a variety of oncogenes. We show that AKT phosphorylation and mTOR activity via enhanced PI3K activity in transformed cells results in enhanced BTN2A1 surface expression, the ligand for the gamma chain of Vγ9Vδ2TCR^14,30^, and thereby makes early transformed cells susceptible for Vγ9Vδ2TCR recognition. However, this step alone is not sufficient for recognition by Vγ9Vδ2TCR T cells, and requires additional upregulation of BTN3A1. We found that in primary pre-cancerous lesions, cells can express either BTN2A1 alone, or BTN2A1 together with additional molecules of the BTN3 complex, which implies earlier tumor control by Vγ9Vδ2T cells in some, but not all pre-cancerous lesions. This observation supports our implication that full activation of T cells via the Vγ9Vδ2TCR requires additional parallel steps. These additional steps include increased levels of intracellular phospho-antigens (pAg), which can be mimicked by aminobisphosphonates, like PAM. Although we could not characterize the natural trigger of BTN3A upregulation, the PAM-model system implied that phosphorylation of the juxtamembrane (JTM) region of BTN3A1 is essential in achieving full recognition by Vγ9Vδ2T cells, and provided a group of kinases most likely responsible for this step. Together, these molecular events result in a tightly regulated composition of membrane expressed BTN2A1 and BTN3A1, which is fed by a constant intracellular protein pool. While intracellular pAg-induced changes in total expression of BTN proteins were minimal, we found additional series of spatial rearrangements directly related to BTN3A1, and identified three novel functionally relevant proteins, CARMIL1, SYNJ2, and PHLDB2, that showed PAM-dependent spatial dynamics.

Our data, in line with previous findings^11,13,30^ imply that BTN2A1 and BTN3A1 membrane expression dynamics are tightly regulated together in human cells. It has been previously suggested that various pathways including AMPK^19^ or EBV-induced activation of JNK^32^ can influence BTN surface expression by enhancing transcription of BTN2A1 and BTN3A. To the contrary, we found the total BTN2A1 protein pool does not differ between normal (healthy) and transformed cells, however, in normal cells we found either no, or low surface expression. And despite the contribution of these studies to an increased understanding of regulation of Vγ9Vδ2T cell targeting, it is still unknown at exactly what stage of tumorigenesis Vγ9Vδ2T cells begin to recognize and eliminate transformed cells. In this study, we show that single mutations that induce higher PI3K activity were sufficient to allow expression of BTN2A1 at the cell membrane, and enable sustained binding of the gamma chain of the Vγ9Vδ2TCR. This observation implies that Vγ9Vδ2TCR-based therapies ^10^ are targeting tumors with oncogenic mutations promoting PI3K activity even with low mutational load. The fact that not only artificial organoids mutated for all tumor associated genes, but also patient derived colorectal organoids are recognized by Vγ9Vδ2TCR cells, shows that these molecular rearrangements are preserved during tumor development, and will allow for targeting of primary, and most likely, metastatic lesions as well. Our data demonstrating that depletion of PI3 kinase activity, (in particular AKT phosphorylation and mTOR activity), in tumor cells impairs Vγ9Vδ2TCR reactivity, implies that targeted therapies acting on the PI3K pathway should be carefully designed, as they might interfere with endogenous innate responses against the tumor. On the other hand, PI3K activation in tumors, next to PAM treatment, might provide a new avenue to possibly sensitize tumors for Vγ9Vδ2TCR-driven attack. Combining the knowledge from these studies with next generation engineering strategies such as TEGs^20,22^, and Gamma delta TCR anti-CD3 bispecific molecules (GABs)^12^ may provide the opportunity to to maximize the targeting potential via Vγ9Vδ2TCRs.

In contrast to the previously mentioned JNK pathway, which is a downstream signaling pathway of PI3K/AKT^33^, AMPK and AKT are often seen to act as antagonists in regulating autophagy and apoptosis^34,35^. On the other hand, under certain conditions, e.g. the presence of cancer, activation of AMPK, especially via the AMPK-activator AICAR, can also activate AKT^36,36^ and mTORC2^37^, and conversely, can also inhibit AMPK via compound C reduced AKT activity^36^. It is also possible that for the initial trigger, BTN2A1 surface expression, an oncogenic PI3K-activating mutation is necessary, but that further regulation and enhancement can be achieved by other pathways, such as AMPK.

While PI3K activity directly affected re-localization of BTN2A1 to the cell membrane, BTN3A1 surface expression remained unaffected by this oncogenic offset. Although we have not been able to identify the precise molecular steps needed for upregulation of BTNs, we show that the ability to be susceptible to Vγ9Vδ2T cells is imprinted early in pre-cancerous lesions. We show that accumulation of intracellular pAg is required, and that dysregulation of the mevalonate pathway is a frequent hallmark of malignant transformation^38^, and that it leads to this accumulation of pAgs. We used aminobisphosphonates (ABP), like PAM as a model system to characterize the next molecular steps. Therefore, the oncogenic-regulated translocation of BTN2A1 to the cell surface and the subsequent binding of pAg to the B30.2 domain of BTN3A1 upon ABP treatment, eventually leads to heterodimerization of BTN2A1 and BTN3A1, and ultimately to TCR binding and activation.

Recently, pAgs were shown to act as the glue between BTN molecules BTN2A1 and BTN3A1 to activate Vγ9Vδ2T cells^18^, and previously, the juxtamembrane (JTM) region was identified as being sensitive for translating pAg-induced inside-out-signaling^17,39^. We have now identified two amino acids that are likely to be phosphorylated specifically upon PAM treatment, and phosphorylation of these sites was associated with higher IFNγ production by Vγ9Vδ2TCR T cells upon co-culture.

Furthermore, our data suggests that phosphorylation of JTM BTN3A1 sites promotes interaction with RhoB and heterodimerization of BTN3A1 and BTN2A1, which will ultimately stabilize BTN2A1 at the cell surface, and will further increase in membrane surface clusters of BTNs that together will form immunological synapses that enable proper T cell activation. These events are not only present in very early transformation events, but are likely also preserved in the late stages of tumors, since colorectal organoids mutated for all four colon cancer associated oncogenes are equally recognized by Vγ9Vδ2TCR T cells as single *APC* mutated.

A closer look at downstream mechanisms after expression of BTN2A1 and BTN3A1 at the cell membrane surface allowed us to reveal that BTN2A1/BTN3A1 surface dynamics are tightly regulated by spatial rearrangements of intracellular modulators which have not yet been described, but are functionally essential to allow Vγ9Vδ2TCR targeting. In this study we established a detailed ABP-dependent interactome of BTN3A1-related proteins using various protein proximity techniques that resolved spatial interaction hierarchies among the studied proteins. PHLDB2 was found, as the only protein of all of the candidates to occupy BTN3A1-proximity membrane clusters, without showing detectable direct close interaction with BTN3A1, implying its role in tissues when no functional recognition by Vγ9Vδ2T cells can yet occur. PHLDB2 dissociates from these clusters either once PAM is present, or in tumor lesions with increased pAg levels, suggesting a complementary role in preventing the accumulation of BTN protein membrane clusters, as also suggested by others^40,41^ **(Figure 5B)**.

We have shown in the past, that active RhoB relocates to the cell membrane in the vicinity of BTN3A1 dimers, and plays a crucial role in the high turn-over of BTN3A1 in cancer cells^16^. Our new data imply that active RhoB might cause re-localization of SYNJ2 from the cytosol to the cell membrane upon PAM stimulation, and might support Rac1, another small GTPase closely linked to RhoB^42^. Our data let us hypothesize that SYNJ2, being enriched in larger size clusters but also in direct interaction with BTN3A1 in the presence of PAM, could lead to a PAM-induced protein aggregation close to BTN3A1. The phosphoinositode-5 phosphatase SYNJ2 has been shown to be involved in actin-based cytoskeleton dynamics^43,44^, and has also been implicated in vesicle trafficking^45^, emphasizing a role in supporting RhoB-mediated BTN3A1 recycling^11^ and enabling recognition of tumor cells by Vγ9Vδ2T cells.

Like SYNJ2 and PHLDB2, CARMIL1 most likely accommodates the formation of cytoskeletal rearrangement^46^ and might altogether be the final step to stabilize BTN2A1-BTN3A1 dimers in the immunological synapse on the tumor cell surface. While we found proof on multiple levels for co-localization or interaction with BTN3A1 in a PAM-dependent manner with CARMIL1, SYNJ2 and PHLDB2, their relationship to BTN2A1 remains unclear. Like AKT1, PHLDB2 contains a PH-domain, which has a high affinity to membrane-bound PIP3^40^, an oncogenic product of PI3K^47^. Therefore, a role of PHLDB2 in recruitment of BTN proteins to the cell membrane is not excluded, and needs to be investigated further.

In summary, and as represented on **Figure 6**, we show that during early mutagenesis, hallmarks for the recognition of a cell through Vγ9Vδ2TCR are induced, such as BTN2A1 surface upregulation and RhoB relocalization. We found that an activated PI3K pathway in single oncogene mutated tumors creates the basis for the susceptibility of cancer cells to Vγ9Vδ2TCR. However, this step is not sufficient for recognition. Increased levels of pAgs, either induced by PAM or endogenously increased as often observed in cancer cells, are essential for recognition by Vγ9Vδ2TCR. Within this context we identified phosphorylation of the BTN3A1 JTM region as an additional independent mechanism needed to achieve a complex recognized by a Vγ9Vδ2TCR. We identified three novel key players, namely CARMIL1, SYNJ2 and PHLDB2, that directly regulate BTN3A1 surface expression, and therefore control BTN2A1-BTN3A1 dimer dynamics on the cell surface. These findings not only shed light on the role of Vγ9Vδ2T cells during early cancer immune surveillance, but also have implications for all Vγ9Vδ2T cell based immune therapies.

## METHODS

### Antibodies

The following antibodies were used: Anti-CD277/BTN3A Alexa Fluor 647 (FAB7316R, Clone 849203), Anti-CD277/BTN3A PE MAb (FAB7316P, Clone 849203), anti-CD277/BTN3A (clone 20.1, LSC106569), pan-γδTCR PE (IMMU510, B49176), Granzyme B APC (QA16A02, 372204), CD107a PE (H4A3), CD8a PerCP-Cy5.5 (RPA-T8, 301032), anti-RhoB mouse monoclonal (C-5, sc-8048), anti-RhoB rabbit polyclonal (abcam, ab170611), anti-CD3 (clone: OKT3), Goat-anti-Rabbit AF488 IgG (H+L), anti-Rabbit AF488 Fab fragment, Goat-anti-Mouse AF488 IgG (H+L), anti-LRRC16A/CARMIL1 Rabbit polyclonal IgG (NBP1-91221), anti-PHLDB2 Rabbit polyclonal IgG (NBP2-38238), anti-CASKIN1 Rabbit polyclonal IgG (HPA055990), anti-HNRNPC mouse monoclonal IgG (AMAb91010), anti-PKP2 Rabbit polyclonal IgG (PA553144), anti-ANKRD26 Rabbit polyclonal IgG (PA559240), anti-VPS13A Rabbit polyclonal IgG (PA554483), anti-TANC1 Rabbit polyclonal IgG (PA557797), anti-SDK2 Rabbit polyclonal IgG (ABIN1386438), anti-NBEA rabbit Polyclonal IgG (HPA040385), anti-PA2G4 Mouse monoclonal IgG (ABIN518579), anti-SYNJ2 Rabbit polyclonal IgG (PA556784). Rabbit mAb ERBB2 (29D8) (1:1,000; #2165; Cell Signaling Technology), rabbit mAb AKT (1:1,000; #9272; Cell Signaling Technology), rabbit Phospho-AKT Ser473 (1:1,000; #9272; Cell Signaling Technology), rabbit mAb Phospho-AKT Thr308 (D25E6) (1:1,000; #13038; Cell Signaling Technology), Rabbit mAb P44/42 MAPK (Erk 1/2) (137F5) (1:1,000; #4695; Cell Signaling Technology), rabbit mAb Phospho-MAPK (Erk 1/2 Thr202/Tyr204) (D13.14.4E) (1:1,000; #4370; Cell Signaling Technology, mouse mAb GAPDH (1;5000, G8795, MERCK/Sigma-Aldrich).

### Cell and patient derived organoid culture

HEK293FT, MDA-MB-231, MZ1851rc, HT29, SKBR-3, Caco-2, HL-60, Phoenix-ampho and MCF10a cells lines were cultured in DMEM+GlutaMAX with 10% fetal calf serum (FCS) and 1% Penicillin-Streptomycin. Daudi cells were cultured in in RPMI-GlutaMAX with 1% Pen/Strep and 10% FCS. MCF10A cells were cultured in DMEM/Ham’s F-12 (Gibco) supplemented with, 1% Penicillin/Streptomycin (Gibco), Ala-Gln (Ultra-Glutamine) (Gibco), 5% heat-inactivated horse serum (Lonza), 0.01 mg/ml insulin (Gibco), 500 ng/ml hydrocortisone (Sigma), 100ng/ml cholera toxin (Sigma), 20 ng/ml epidermal growth factor (EGF) (Peprotech). Cell lines were routinely tested for mycoplasma and STR type verified by eurofins. PBMCs were isolated from buffy coats using ficoll-paque obtained from the Sanquin blood bank. Colon organoids were established and cultured as previously described^23,48^. Before co-cultures, organoids were recovered from the BME using TrypLE express.

### RNA extraction and library preparation

RNA was extracted using standard protocols and quantified to ensure high integrity. Libraries were prepared using the Illumina RNA library preparation kit, following the manufacturer’s instructions. Libraries were sequenced on an Illumina NextSeq 500 platform in paired-end mode (76:i7, 8:i5). Reads were trimmed for quality and adapters and aligned to the human genome (*GRCh38*) using *STAR* with default parameters. Gene-level counts were generated using *featureCounts*.

### Generation of engineered T-cells

γδTCR and HER2 CAR T-cells were generated as previously published^22^. In short, Phoenix-ampho cells were transfected with env (COLT-GALV), gag-pol (HIT60) and pBullet retrovirus constructs containing either both TCR chains or the HER2 CAR sequence. Pre-activated T-cells were subsequently transduced twice, with the viral supernatant of these cells with polybrene. Transduced T-cells were isolated and expanded using a rapid expansion protocol.

### Viral transduction MCF10a

pBABE-Puro retroviral vector (EV) and pBABE-Puro vectors carrying ERBB2 wild-type and mutants were co-transfected independently with pUMVC (Addgene #8449) and VSV-G (Addgene #8454) retroviral packaging plasmids into HEK-293T cells using PEI-transfection. Medium was refreshed after 24hrs and viral supernatant was collected 48hrs and 72hrs after transfection. Viral particles were added to low passage MCF10A cells and incubated overnight. Twenty-four hours following transduction, MCF10A cells were selected using 2 µg/ml Puromycin (Gibco) for two days. Gene transduction and protein expression were validated using Western analysis. Established cell lines were immediately expanded and cryopreserved in low passage aliquots.

### Plasmids

Retroviral pBabe-Puro plasmids were purchased from Addgene: ERBB2 (#40978), ERBB2::S310F (#40991), and ERBB2:: A775_G775insV, G776C (V777E) (#40979). All plasmids and mutations were verified by Sanger Sequencing.

### BioID

pcDNA3.1 MCS-BirA(R118G)-HA was a gift from Kyle Roux (Addgene plasmid # 36047; http://n2t.net/addgene:36047; RRID:Addgene_36047). The DNA sequence of BTN3A1-BirA(R118G)-HA has been subcloned into pBullet retrovirus constructs. Phoenix-ampho cells were transfected with env (COLT-GALV), gag-pol (HIT60) and pBullet retrovirus constructs containing BTN3A1-BirA(R118G)-HA constructs. HEK293FT cells were subsequently transduced twice with the viral supernatant of these cells with polybrene. Transduced cells were selected using 2,5 mg/ml G418. Expression was confirmed by confocal microscopy. The preparation of the samples for analysis by LS-MS followed the protocol by Roux et al ^31,49^. In short, HEK293FT WT or BTN3A1-BirA(R118G)-HA expressing cells were 50 µM Biotin alone, or 50 µM Biotin and 100 µM Pamidronate. Cells were lysed and sonicated to generate whole cell lysates. Biotinylated proteins were pulled down using streptavidin-magnetic beads, and samples were then analyzed by LS-MS. Each condition was prepared as quadruplicate.

### Western Blot

Cell lines were seeded in a 10 cm dish overnight to confluency. Cells were rinsed with ice cold PBS and lysed with lysis buffer containing 1% NP-40, 150 mM NaCl, 20 mM Tris HCl pH 7,6 and protease inhibitors (complete, Roche, #11873580001). Protein concentration was determined using Pierce BCA Protein Assay Kit - Rapid Gold (Fisher Scientific, #15776178) and 5 µg total protein was loaded on each lane of a Mini Protean TGX Gel (BioRad, #4561093) together with a standard (WesternC, BioRad, #1610376) and run at 140V. The protein was then transferred to a 0.2 µm nitrocellulose membrane (BioRad, #1704158) using Trans-Blot Turbo System. The membrane was incubated in blocking buffer (PBS-T + 5% BSA for 1h at RT, washed three times with PBS-T (0,1% Tween) and incubated overnight on a rotator with the respective primary antibodies as indicated. After washing three times with PBS-T at RT, membranes were incubated with the respective secondary antibodies for 1h at RT on a rotator. After final washings, membranes were developed using the ECL Kit and were measured.

### Inhibitors

The following inhibitors were used in this study; PI3K inhibitor GDC-0941 (Pictilisib) (SelleckChem, #S1065), pan-AKT inhibitor MK2206 (SelleckChem, #S1078), MEK inhibitor AZD6244 (Selumetinib) (SelleckChem, #S1008), mTORC 1 inhibitor Rapamycin (Sirolimus) (SelleckChem, #S1039), mTORC 1 / 2 inhibitor Torin 1 (SelleckChem, #S2827).

### Stimulation assays

50.000 tumor cells were co-cultured overnight at 37°C in a round-bottom 96-well plate (Nunc, Thermo-Fisher Scientific) with 50.000 Vγ9Vδ2-expressing T cells (TEGs) or HER2 CAR expressing T cells in RPMI-GlutaMAX medium, containing pamidronate as indicated. Furthermore, for the inhibitor experiments, tumor cells were pre-treated overnight with either PI3K (Pictilisib, GDC-0941), AKT (MK2206), MEK (Selumetinib, AZD6244) inhibitors, Rapamycin or Torin. Before co-culture, cells were washed twice with fresh RPMI medium, and afterwards were adjusted to 50.000 cells in 100 µL RPMI culture medium. After co-culture, 100µL supernatant was used for INF-γ detection using the Invitrogen^TM^ eBioscience^TM^ human IFN-γ ELISA kit ready-set-go by following the manufacturer’s protocol^42^.

### Flow Cytometry

Cells were counted and 200.000 per condition were put in a FACS tube, after which 1mL FACS buffer (PBS, 1% Na-azide) was added and tubes spun down at 1500RPM for 5 min. Supernatant was discarded and, for the tetramer and bead stainings with soluble TCRs, cells were incubated with tetramer/bead solution for 30 minutes at RT. Cells are washed with PBS and subsequently, 20µL of the antibody mixture in FACS buffer was added to each well, after which cells were resuspended and incubated for 30min at RT. 1mL FACS buffer was added to each well, and cells were spun down at 1500RPM for 5 min, supernatant was removed and resuspended in 1mL FACS buffer and spun down again at 1500RPM for 5 min. Finally, supernatant was removed and 150µL 1% PFA was added. Samples were measured on a FACSCanto 2 (BD bioscience) using FACSDiva software.

### Intracellular staining and colocalization analysis

MZ1851rc cells were seeded in a 16-well Nunc Lab-Tek^TM^ chamber at 1500 cells per well in 200µL cDMEM, and cultured for 76h at 37°C, 5% CO2 in a wet chamber (petri dish + wet paper cloth). Medium was removed and 200µL cDMEM with 100µM pamidronate (PAM) (where indicated) was added to the wells. After 24h incubation, cells were washed with PBS and subsequently fixed with 4% paraformaldehyde (PFA) for 15min at RT. Cells were permeabilized for 10min using either 100% ice-cold methanol, 0.3% Triton X100 in PBS, or 0.1% saponin in PBS (changed protocol over time, saponin seems to work best) at RT. Sample blocking was performed by incubating wells for 1h at RT with blocking solution (PBS, 5% Pooled human serum (HuS), 1% BSA, 1% Na-azide + either 0.3% Triton X-100 or 0.1% saponin). Primary antibodies diluted in antibody dilution buffer (PBS, 1% BSA, 1% Na-azide + either 0.3% Triton X-100 or 0.1% saponin) were added to the wells and were incubated either 1h at RT, or overnight at 4°C. Cells were washed 3x with PBS and a secondary antibody diluted in antibody dilution buffer was added and incubated either 1h at RT, or overnight at 4°C. Finally, the chamber gasket and any glue residue was removed from the well chamber using a sterile tweezer and surgical blade, some Prolong Diamond Antifade mountant with DAPI was added and a coverglass applied. Before imaging, any leftover residue was removed with a Kimwipe and the slide was kept in a dark place for at least 30min. Imaging was performed on a Zeiss LSM710 confocal laser scanning microscope using a 63x 1.40 oil immersion objective. Analysis of pictures was done blinded to conditions using Volocity^TM^ image analysis software (PerkinElmer) by selecting areas of interest on membrane clusters of BTN3A, and determining the Pearson’s correlation coefficient. Representative images were made using ImageJ software (NiH).

### Animal models

NSG mice were administered total body irradiation of 1.65 Gy on day -1. AKPS mutated PDOs were injected subcutaneously in the right flank on day 0. On day 1 and 7, 10^7^ TEG-001 or TEG-LM1 cells were administered intravenously in pamidronate (10 mg/kg body weight). Next to T-cell administration, 0.6 × 106 IU of IL-2 in incomplete Freund’s adjuvant was injected subcutaneously at day 1 as described in^50^. Bioluminescence was used to weekly monitor PDO outgrowth over time.

### Proximity ligation assays

Duolink^TM^ PLA Fluorescence protocol was followed.^41^ Duolink Plus and minus probes and detection reagents orange were obtained from Merck. MZ1851rc cells were seeded in a 16-well Nunc Lab-Tek^TM^ chamber (Thermo-Fisher Scientific, Nunc) at 15000 per well per 200µL cDMEM and cultured for 76h. Afterwards, medium was gently removed and 100µM pamidronate in cDMEM was added where indicated, and incubated overnight. Wells were washed 1x with PBS and fixed with 4% PFA for 15min at RT. Wells were washed 3x with PBS and permeabilized using 0.1% saponin in PBS for 15min at RT. Wells were blocked using Duolink blocking buffer and primary antibodies diluted in Duolink PLA diluent were added to the wells, and incubated for 1h at RT in a humidity chamber (empty pipette tip box with elevated plateau covered in parafilm for slide to sit on, water around it). Wells were washed 2x with 1x wash buffer A (Merck) and incubated for 1 hour at 37°C in a wet chamber with 35µL/well of Duolink min/plus PLA probe mix (anti-mouse-plus, anti-rabbit-minus). Wells were washed 2x with 1x wash buffer A and incubated with 35µL/well Duolink ligation buffer with 1:40 Ligase enzyme added from a freezing block and incubated for 30 min at 37°C in a wet chamber. Wells were washed 2x with 1x wash buffer A again and incubated with Duolink Orange amplification buffer with 1:80 polymerase enzyme added from a freezing block and incubated for 100min at 37°C. Finally, wells were washed 2x with 1x wash buffer B (Merck) for 10min and 1x with 0.01x wash buffer B for 1 min. All remaining buffer was discarded and further preparation of the slide was done in the same way as described for intracellular staining. Imaging was performed on a Zeiss LSM710 confocal laser scanning microscope using both a 20x objective or a 63x 1.40 oil immersion objective. PLA clusters per image and number of nuclei were determined using Volocity^TM^ image analysis software (PelkinElmer).

### Fluorescence Resonance Energy Transfer (FRET)

To study direct interaction between proteins upon phosphoantigen accumulation, HEK293T cells expressing phospho-variants of BTN3A1 were first dissociated using Trypsin-EDTA, transferred into a FACS tube and resuspended in complete DMEM. The cells were then treated with 100 μM PAM for 1 hour at 37°C and washed with FACS buffer (PBS, 1% Na-azide). After that, the cells were blocked with PBS containing 5% BSA for 15 minutes and then incubated for 30 minutes at room temperature, with an antibody conjugated with the donor fluorochrome at 100μl staining volume. Cells were washed once and then split into two samples: donor and donor+acceptor. Donor samples were incubated with FACS buffer and donor+acceptor samples were incubated for 30 minutes at room temperature, with an antibody conjugated with the acceptor fluorochrome at 50μl staining volume. After washing with FACS buffer twice, samples were fixed with 1% PFA. The donor fluorescence was measured using a FACS Canto-II flow cytometer (BD Biosciences) where donor fluorescence of the donor+acceptor (double stained) samples was compared with one of the samples labeled only with the donor antibody. FRET efficiency was calculated from the fractional decrease of the donor fluorescence in the presence of the acceptor. Background noise in the donor fluorescent channel due to spectral overlap with different fluorescence channels was excluded from the calculations, by subtracting the measured mean fluorescence intensity (MFI) on unlabeled and single stained samples from the MFI of the donor and donor+acceptor samples respectively. The centrifugation steps during washes were done at 1500RPM for 5 minutes, and the antibodies were diluted in FACS buffer. The samples stained with antibodies conjugated to Alexa Fluor 594 were validated for proper staining by remeasurement using an LSR Fortessa cell analyzer (BD Biosciences).

### Generation of HEK-293FT phospho-variants

For the generation of the cell lines stably expressing BTN3A1 phosphovariants and BTN3A2, the BTN3A1-Flag as well as the BTN3A2 sequence were codon optimized, custom synthesized and cloned in pBullet-IRES-puro plasmid vectors. The retroviral particles produced by transfection in Phoenix ampho cells were generated with the same procedure used to generate TEGs. The viral supernatant from the Phoenix ampho cells was used to transduce the HEK 293T BTN3KO cell line. 48 hours post-transduction 1.5 μg/mL of puromycin was supplemented to the culture medium for antibiotic selection and it was carried out until the control (untransduced) cell line died completely.

### RNA-sequencing data analysis

The downstream analysis was performed in R mainly using the following packages: *DESeq2*, *ggplot2*, *ComplexHeatmap*, GSVA, and *AnnotationDbi* with *org.Hs.eg.db* for annotation. Preprocessing steps included normalization, filtering low-count genes, and variance stabilizing transformation. Gene set variation analysis (GSVA) was performed to assess pathway enrichment using hallmark gene sets from MSigDB. No new software or tools were generated during the analysis. Raw RNA-seq data will be made available through the European Genome-Phenome Archive (EGA).

### Software used

Office 2016 (Microsoft), Illustrator CS6 / Illustrator 2019 (Adobe systems), Zen 2009 / 2012 (Zeiss), ImageJ (NiH), Volocity image analysis software (PerkinElmer), GraphPad Prism 8, Microplate Manager (Bio-Rad), FACSDiva (BD Biosciences), R software (R Core Team, version 4.1.2).

## Acknowledgements

We thank the staff of the Flow Core Facility at the UMC Utrecht. We kindly thank Prof. Erin Adams (The University of Chicago) for providing the CD277 KO HEK293T cell line and Halvard Boenig (Institute for Transfusion Medicine and Immunohematology, Goethe University, Frankfurt a. M., Germany) for providing feeder cells. Funding for this study was provided by ZonMW 43400003 and VIDI-ZonMW 917.11.337, UU 2013-6426, UU 2014-6790 and UU 2015-7601 and Gadeta to J.K., UU 2018-11393 to Z.S. and J.K., Marie Curie 749010 to D.X.B. D.F., W.W., M.A. and A.J.R.H. We acknowledge financial support from the NWO funded Netherlands Proteomics Centre through the National Road Map for Large-scale Infrastructures program X-Omics (Project 184.034.019).

## Contributions

A.C., A.D.M., D.F., T.K., P.B., I.J., D.X.B., P.H.L., S.H., L.G., T.M., S.L., M.H., J.B., D.V.D., E.S.J., J.S., N.T., L.H.H., A.N. and T.A. performed the experiments and analyzed data, W.W., T.S., H.S., J.R., P.D., J.D., A.J.R.H., H.C., J.K. and Z.S. supervised the study, J.K. and Z.S. designed the study, A.C. and A.D.M. wrote the paper together with Z.S. and J.K.

