## Supplemental Figures for "Sensitivity to Vγ9Vδ2TCR T cells is imprinted after single mutations during early oncogenesis"

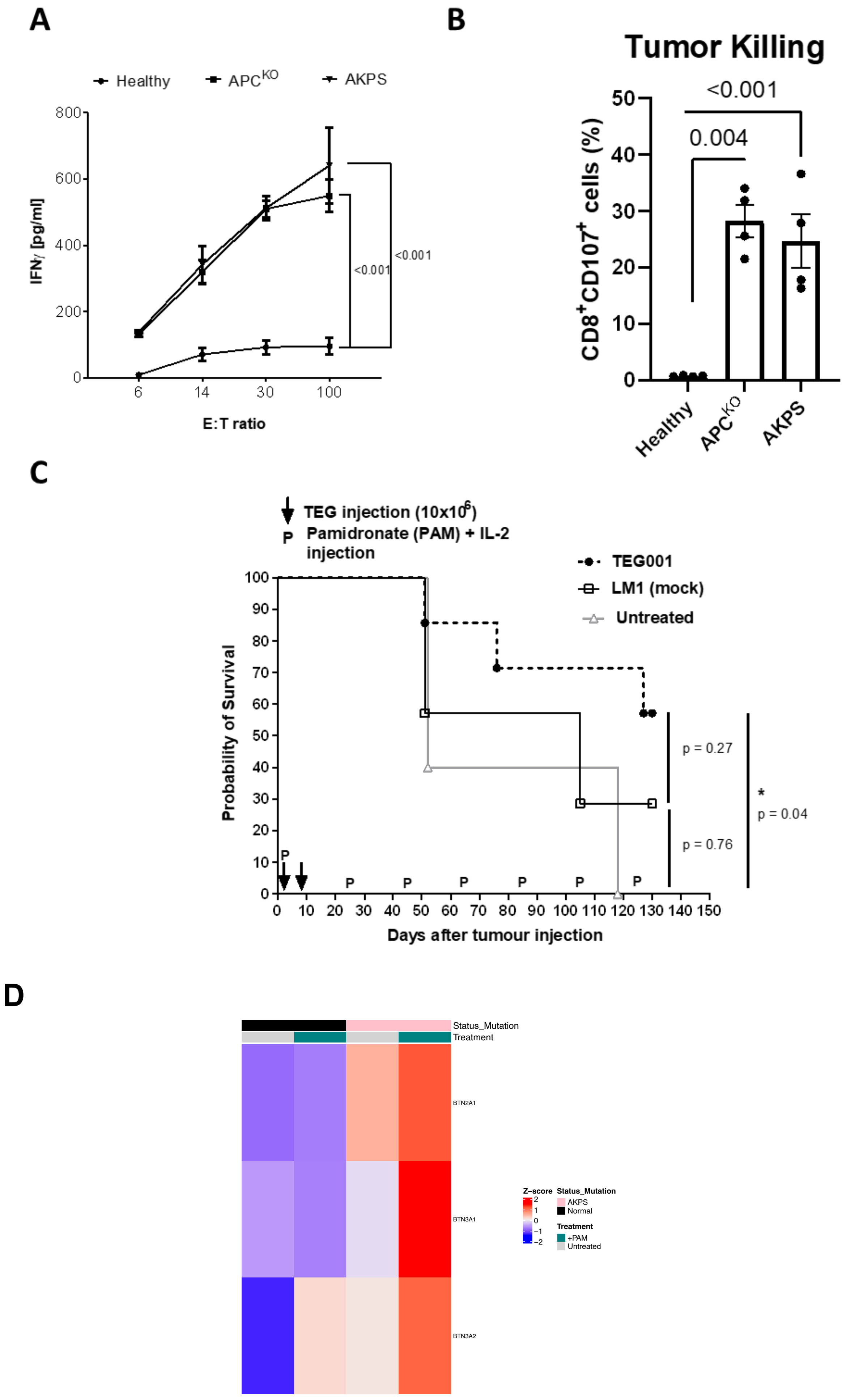
A
Healthy
APC^KO
AKPS
800
600
400
200
0
IFNγ [pg/ml]
6
14
30
100
E:T ratio
<0.001
<0.001
B
Tumor Killing
<0.001
0.004
50
40
30
20
10
0
CD8+CD107+ cells (%)
Healthy
APC^KO
AKPS
C
TEG injection (10x10^6)
P Pamidronate (PAM) + IL-2 injection
TEG001
LM1 (mock)
Untreated
Probability of Survival
100
90
80
70
60
50
40
30
20
10
0
P
p = 0.27
p = 0.76
*
p = 0.04
0 10 20 30 40 50 60 70 80 90 100 110 120 130 140 150
Days after tumour injection
D
Status_Mutation
Treatment
BTN2A1
BTN3A1
BTN3A2
Z-score
2
1
0
−1
−2
Status_Mutation
AKPS
Normal
Treatment
+PAM
Untreated

**E**

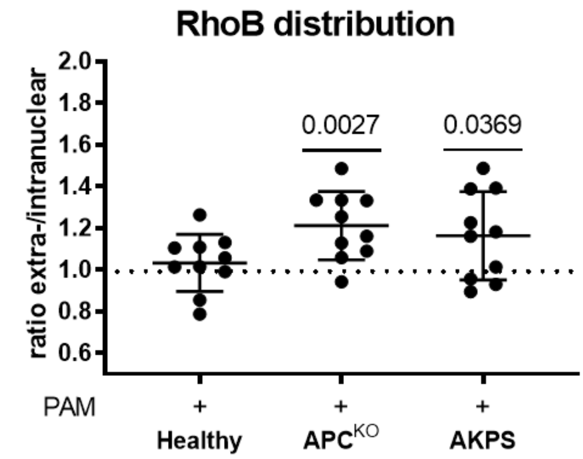


**Supplementary Figure 1. TEG targeting of APC and AKPS. (A)** IFNγ production by Vγ9Vδ2TCR T cells after co-culture with wt, APC and APKS CRC organoids at various target:effector ratios in the presence of 100uM PAM. **(B)** CD107 degranulation of Vγ9Vδ2TCR T cells after co-culture with wt, APC and APKS CRC organoids at various PAM concentrations. **(C)** NOD.Cg-Prkdcscid Il2rgtm1Wjl/SzJ (NSG) mice were injected subcutaneously in right-flank with 250,000 cells AKPS CRC organoid (suspended in culture media and Matrigel with 1:1 ratio) on Day 0 followed by intravenous injection of either PBS (untreated control), $10^7$ TEG001 or TEG-LM1 cells on Day 1 and 6 (n =7 mice; indicated by arrows). Overall survival of tumor-bearing mice was recorded for 130 days. Statistical significance was calculated by log-rank (Mantel-Cox) test, * P <0,05. **(D) Expression of BTNx genes in the WT-AKPS model with and without PAM (presented as z-scores normalized by gene) (E)** RhoB intracellular distribution after PAM treatment in wt, $APC^{KO}$ or AKPS mutant CRC organoids, measured by confocal microscopy.

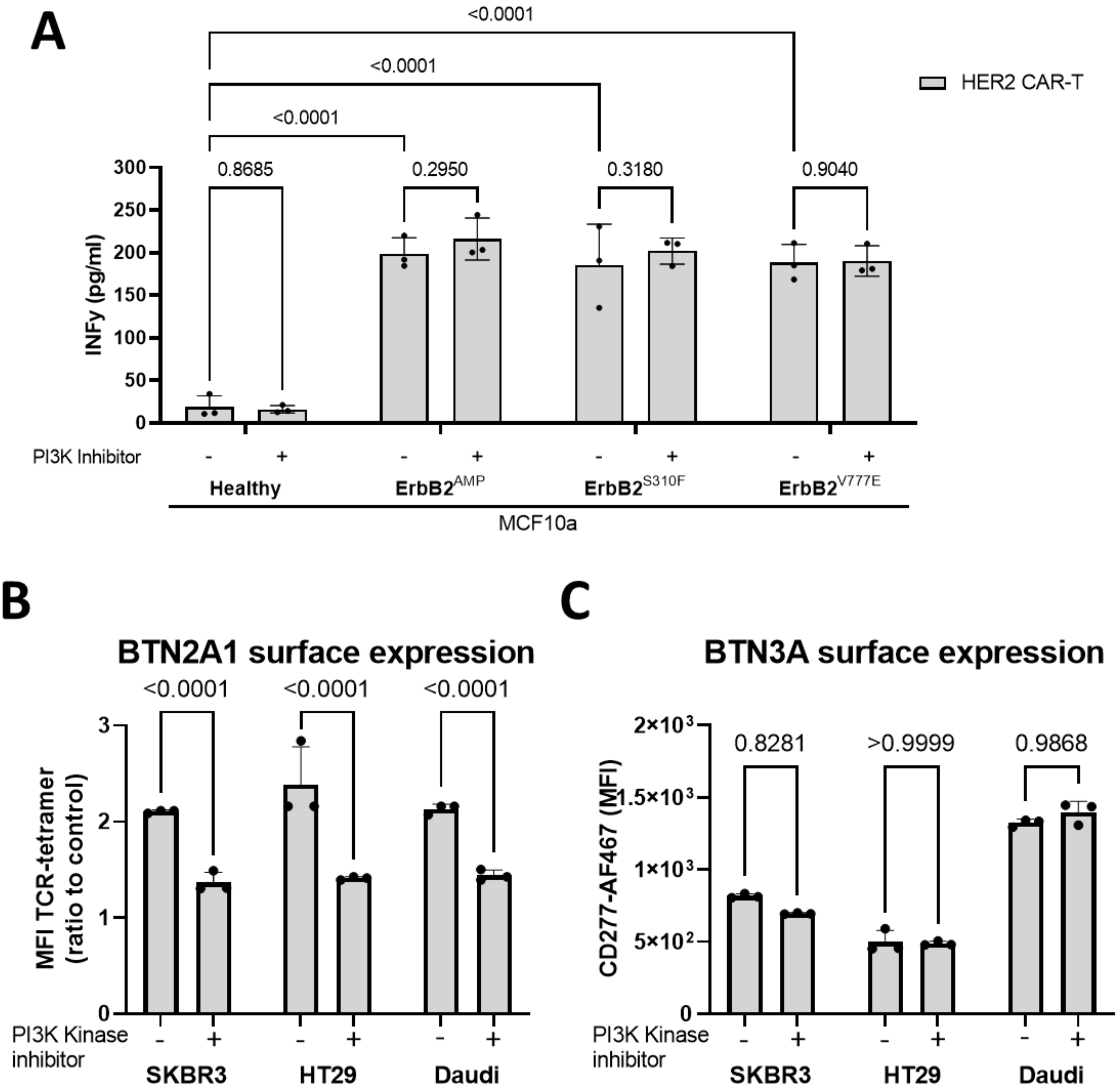


**Supplementary Figure 2. Addition of PI3K inhibitor consistently reduces BTN2A1 surface expression, but not BTN3A surface expression, in various tumor cell lines. (A)** MCF10a wt (healthy) and ErbB2/HER2 mutants were co-cultured overnight with HER2 CAR-T cells. IFNγ expression was evaluated with IFNγ ELISA. **(B)** BTN2A1 surface expression was determined by Vγ9Vδ2TCR-tetramer binding relative to control-tetramer binding after pre-incubation with PI3K inhibitor. **(C)** BTN3A surface expression was measured using a CD277-AF467 antibody after pre-incubation with PI3K inhibitor.

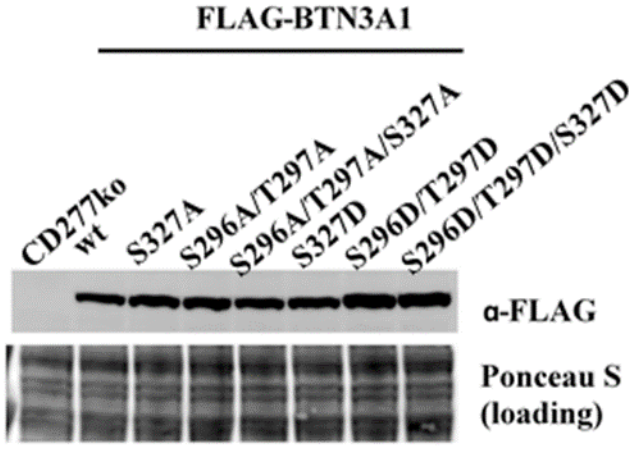


**Supplementary Figure 3. Total expression of BTN3A1 phosphor-variants.** Western blot analysis of HEK-293 BTN3A1 KO cells reconstituted with either wt BTN3A1 of with phospho-variants of BTN3A1 using anti-FLAG antibody.

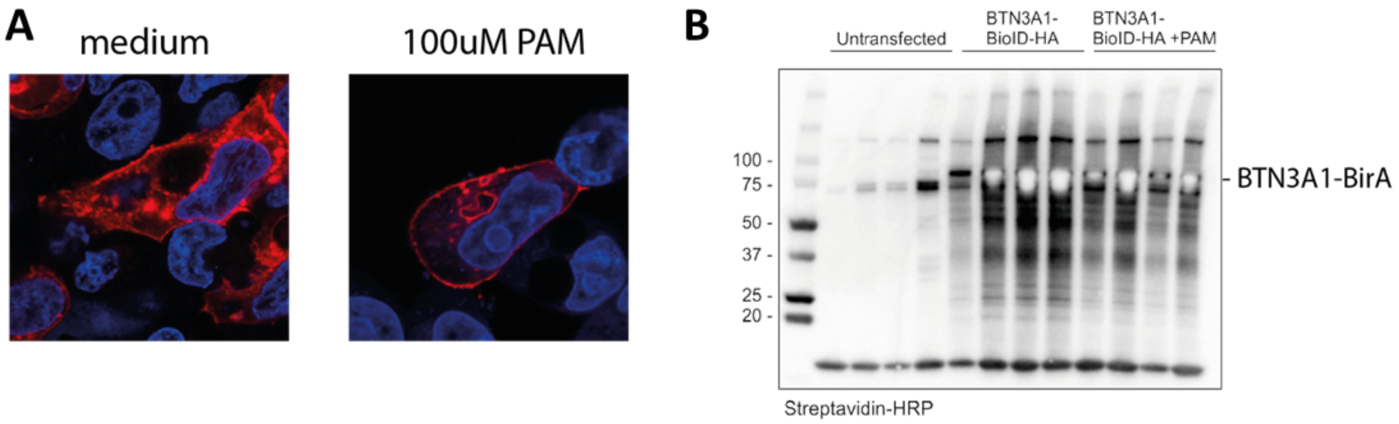


C

BTN3A1-interacting proteins (List provided by Mass Spectrometry, David Vargas Diaz)

| PAM | | | | | |
|---|---|---|---|---|---|
| siRNA Nr. | Sigma Nomenclature siRNA | Gene Name | Full Protein Name | Uniprot ID | Probability of being an specific hit |
| 1 | SASI_Hs01_00246154 | GOLGA3 | Golgin subfamily A member 3 | Q08378 | 0,98 |
| 2 | SASI_Hs02_00323533 | GIGYF2 | GRB10-interacting GYF protein 2 | Q6Y7W6 | 0,98 |
| 3 | SASI_Hs02_00346828 | TNIK | TRAF2 and NCK-interacting protein kinase | Q9UKE5 | 0,93 |
| 4 | SASI_Hs01_00056160 | EPB41L5 | Band 4.1-like protein 5 | Q9HCM4 | 0,86 |
| 5 | SASI_Hs01_00138764 | PKP4 | Plakophilin-4 | Q99569 | 0,78 |
| 6 | SASI_Hs01_00242266 | ARHGAP39 | Rho GTPase-activating protein 39 | Q9C0H5 | 0,76 |
| 7 | SASI_Hs01_00024230 | MINK1 | Misshapen-like kinase 1 | Q8N4C8 | 0,75 |
| 8 | SASI_Hs01_00096844 | EIF2AK3 | Eukaryotic translation initiation factor 2-alpha kina | Q9NZJ5 | 0,73 |
| 9 | SASI_Hs02_00305254 | IGSF3 | Immunoglobulin superfamily member 3 | O75054 | 0,73 |
| 10 | SASI_Hs01_00175890 | PDZD8 | PDZ domain-containing protein 8 | Q8NEN9 | 0,71 |
| 11 | SASI_Hs02_00362041 | PARD3B | Partitioning defective 3 homolog B | Q8TEW8 | 0,7 |
| 12 | SASI_Hs01_00083047 | STBD1 | Starch-binding domain-containing protein 1 | O95210 | 0,67 |
| 13 | SASI_Hs02_00338274 | KCNB2 | Potassium voltage-gated channel subfamily B mem | Q92953 | 0,66 |
| 14 | SASI_Hs02_00356911 | FAT4 | Protocadherin Fat 4 | Q6V017 | 0,66 |
| 15 | SASI_Hs01_00140637 | PKP2 | Plakophilin-2 | Q99959 | 0,65 |
| 16 | SASI_Hs01_00054582 | SDK2 | Protein sidekick-2 | Q58EX2 | 0,65 |
| 17 | SASI_Hs02_00312564 | HNRNPC | Heterogeneous nuclear ribonucleoproteins C1/C2 | P07910 | 0,65 |
| 18 | SASI_Hs01_00068769 | SYNJ2 | Synaptojanin 2 | O15056 | 0,63 |
| 19 | SASI_Hs01_00183050 | LRRC16A | F-actin-uncapping protein LRRC16A/Capping protei | Q5VZK9 | 0,63 |
| 20 | SASI_Hs01_00041017 | CDC42BPA | Serine/threonine-protein kinase MRCK alpha | Q5VT25 | 0,62 |
| 21 | SASI_Hs01_00197680 | VPS13A | Vacuolar protein sorting-associated protein 13A | Q96RL7 | 0,62 |
| 22 | SASI_Hs01_00022791 | PA2G4 | Proliferation-associated protein 2G4 | Q9UQ80 | 0,62 |
| 23 | SASI_Hs02_00361260 | TANC1 | Tetratricopeptide repeat, ankyrin repeat and coiled | Q9C0D5 | 0,62 |
| 24 | SASI_Hs01_00095038 | LIMD1 | LIM domain-containing protein 1 | Q9UGP4 | 0,62 |
| 25 | SASI_Hs01_00156891 | COBLL1 | Cordon-bleu protein-like 1 | Q53SF7 | 0,62 |
| 26 | SASI_Hs02_00333957 | NEO1 | Neogenin | Q92859 | 0,62 |
| 27 | SASI_Hs01_00247261 | SYNCRIP | Heterogeneous nuclear ribonucleoprotein Q/Synap | O60506 | 0,62 |
| 28 | SASI_Hs01_00152399 | OSBPL3 | Oxysterol-binding protein-related protein 3 | Q9H4L5 | 0,62 |
| 29 | SASI_Hs01_00026464 | TPR | Translocated promoter region protein | P12270 | 0,6 |

| No PAM | | | | | |
|---|---|---|---|---|---|
| | Sigma Nomenclature siRNA | Gene Name | | Uniprot ID | Probability of being an specific hit |
| 2 | SASI_Hs02_00323533 | GIGYF2 | | Q6Y7W6 | 0,98 |
| 1 | SASI_Hs01_00246154 | GOLGA3 | | Q08378 | 0,94 |
| 16 | SASI_Hs01_00054582 | SDK2 | | Q58EX2 | 0,88 |
| N1 | SASI_Hs02_00344266 | RAB3GAP2 | | Q9H2M9 | 0,86 |
| 3 | SASI_Hs02_00346828 | TNIK | | Q9UKE5 | 0,84 |
| 4 | SASI_Hs01_00056160 | EPB41L5 | | Q9HCM4 | 0,84 |
| 5 | SASI_Hs01_00138764 | PKP4 | | Q99569 | 0,8 |
| N2 | SASI_Hs02_00310578 | EIF4G2 | | P78344 | 0,75 |
| N3 | SASI_Hs01_00065570 | ROCK1 | | Q13464 | 0,73 |
| N4 | SASI_Hs01_00012614 | SMCR8 | | Q8TEV9 | 0,68 |
| N5 | SASI_Hs01_00197359 | SH3PXD2B | | A1X283 | 0,68 |
| N6 | SASI_Hs02_00350925 | DENND4C | | Q5VZ89 | 0,67 |
| N7 | SASI_Hs02_00307420 | KPRP | | Q5T749 | 0,66 |
| N8 | SASI_Hs02_00315355 | USP6NL | | Q92738 | 0,66 |
| 11 | SASI_Hs02_00362041 | PARD3B | | Q8TEW8 | 0,66 |
| N9 | SASI_Hs01_00227359 | PDE3B | | Q13370 | 0,66 |
| N10 | SASI_Hs01_00183260 | PPP1R9A | | Q9ULJ8 | 0,66 |
| N11 | SASI_Hs01_00112864 | ANKRD26 | | Q9UPS8 | 0,65 |
| N12 | SASI_Hs02_00322254 | CABLES1 | | Q8TDN4 | 0,64 |
| N13 | SASI_Hs02_00353387 | PCDH9 | | Q9HC56 | 0,64 |
| N14 | SASI_Hs01_00086890 | ERC1 | | Q8IUD2 | 0,64 |
| 8 | SASI_Hs01_00096844 | EIF2AK3 | | Q9NZJ5 | 0,64 |
| N15 | SASI_Hs02_00327487 | NCOA7 | | Q8NI08 | 0,64 |
| N16 | SASI_Hs01_00098727 | ADCY9 | | O60503 | 0,64 |
| N17 | SASI_Hs01_00191556 | UNC45A | | Q9H3U1 | 0,64 |
| N18 | SASI_Hs02_00350287 | NOTCH1 | | P46531 | 0,63 |
| N19 | SASI_Hs02_00347838 | CAMSAP1 | | Q5T5Y3 | 0,63 |
| N20 | SASI_Hs01_00138034 | PCDH17 | | O14917 | 0,63 |
| N21 | SASI_Hs01_00031229 | PHLDB2 | | Q86SQ0 | 0,63 |
| N22 | SASI_Hs01_00079280 | CLMN | | Q96JQ2 | 0,63 |
| 10 | SASI_Hs01_00175890 | PDZD8 | | Q8NEN9 | 0,62 |
| N23 | SASI_Hs01_00177000 | PCDHGA11 | | Q9Y5H2 | 0,62 |
| 14 | SASI_Hs02_00356911 | FAT4 | | Q6V017 | 0,62 |
| N24 | SASI_Hs02_00348326 | NBEA | | Q8NFP9 | 0,62 |
| N25 | SASI_Hs01_00178257 | CASKIN1 | | Q8WXD9 | 0,62 |
| N26 | SASI_Hs02_00311332 | TMEM237 | | Q96Q45 | 0,62 |
| N27 | SASI_Hs01_00106398 | PNN | | Q9H307 | 0,62 |
| N28 | SASI_Hs01_00046208 | FAM83B | | Q5T0W9 | 0,61 |
| N29 | SASI_Hs01_00158604 | GOLGB1 | | Q14789 | 0,61 |
| N30 | SASI_Hs02_00313610 | SDK1 | | Q7Z5N4 | 0,61 |
| N31 | SASI_Hs01_00171660 | STK10 | | O94804 | 0,6 |

| |
|---|
| -/+ PAM |
| + PAM |
| -PAM |

**Supplementary Figure 4. Expression of BTN3A1-BirA* of candidate protein. (A)** Representation of differentially expressed genes between AKPS CRC organoids incubated either with or without 100uM PAM. **(A)** HEK-293F cells were transfected with BTN3A1-BirA fusion plasmid and biotinylating pattern of fusion protein was controlled after adding biotin. Biotinylation was visualized by adding streptavidin-APC (red) and DAPI (blue) for nuclear staining. **(B)** cellular expression of the fusion protein was controlled by western blot. **(C)** List of BTN3A1-BioID biotinylated proteins identified by mass spectrometry with a > 0,6 probability being a specific hit. Colours indicate in which condition the proteins were found under various conditions (green = no PAM, blue = PAM, orange = both).

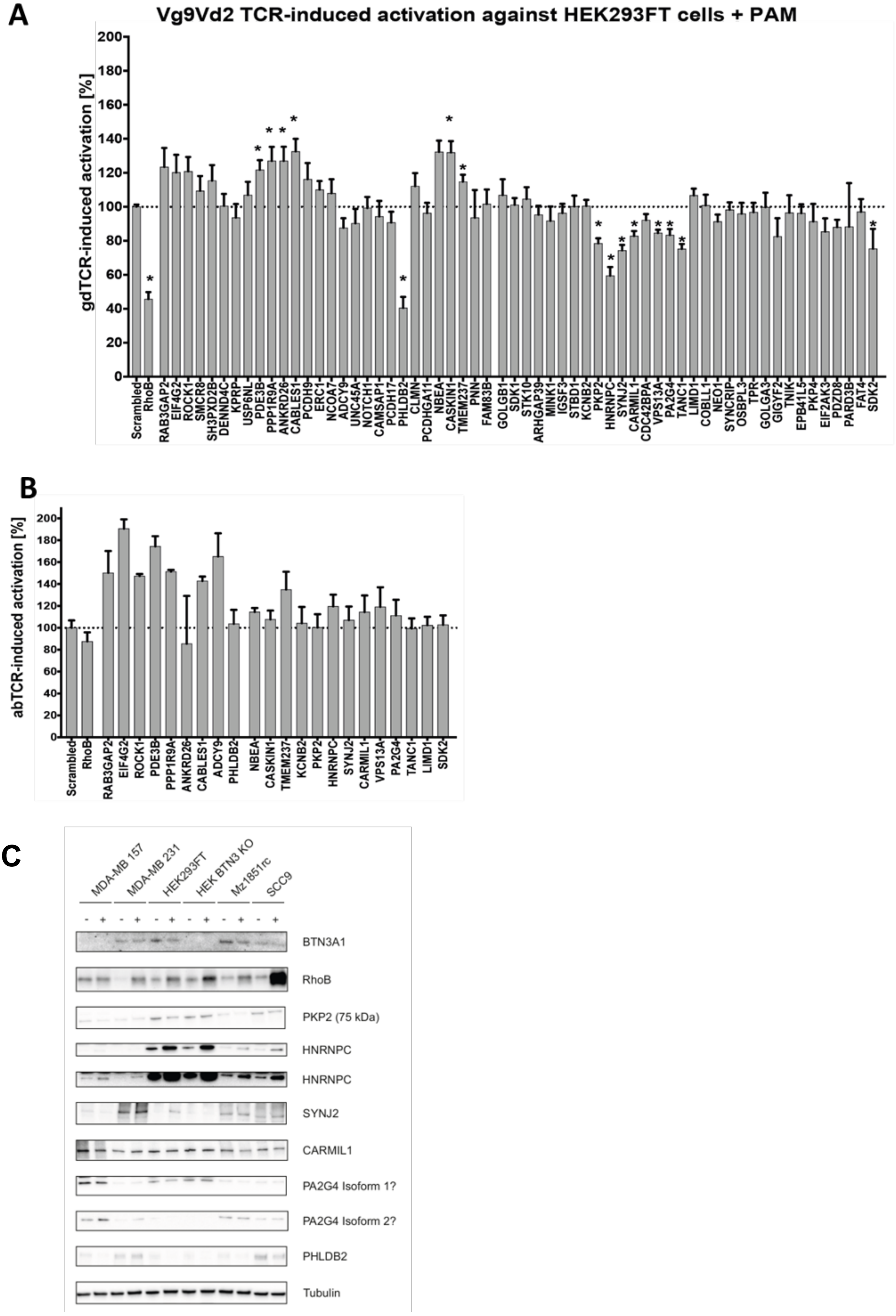
A
Vg9Vd2 TCR-induced activation against HEK293FT cells + PAM
gdTCR-induced activation [%]
0 20 40 60 80 100 120 140 160 180 200
Scrambled RhoB RAB3GAP2 EIF4G2 ROCK1 SMCR8 SH3PXD2B DENND4C KPRP USP6NL PDE3B PPP1R9A ANKRD26 CABLES1 PCDH9 ERC1 NCOA7 ADCY9 UNC45A NOTCH1 CAMSAP1 PCDH17 PHLDB2 CLMN PCDHGA11 NBEA CASKIN1 TMEM237 PNN FAM83B GOLGB1 SDK1 STK10 ARHGAP39 MINK1 IGSF3 STBD1 KCNB2 PKP2 HNRNPC SYNJ2 CARMIL1 CDC42BPA VPS13A PA2G4 TANC1 LIMD1 COBLL1 NEO1 SYNCRIP OSBPL3 TPR GOLGA3 GIGYF2 TNIK EPB41L5 PKP4 EIF2AK3 PDZD8 PARD3B FAT4 SDK2
B
abTCR-induced activation [%]
0 20 40 60 80 100 120 140 160 180 200
Scrambled RhoB RAB3GAP2 EIF4G2 ROCK1 PDE3B PPP1R9A ANKRD26 CABLES1 ADCY9 PHLDB2 NBEA CASKIN1 TMEM237 KCNB2 PKP2 HNRNPC SYNJ2 CARMIL1 VPS13A PA2G4 TANC1 LIMD1 SDK2
C
MDA-MB 157
MDA-MB 231
HEK293FT
HEK BTN3 KO
Mz1851rc
SCC9
- + - + - + - + - + - +
BTN3A1
RhoB
PKP2 (75 kDa)
HNRNPC
HNRNPC
SYNJ2
CARMIL1
PA2G4 Isoform 1?
PA2G4 Isoform 2?
PHLDB2
Tubulin

**Supplementary Figure 5. siRNA screening of protein candidates. (A)** HEK-293FT cells were transfected with siRNAs against candidate proteins and cells were used as targets against Vγ9Vδ2TCR T cells in the presence of 100uM PAM and IFNγ release of T cells upon activations was measured by ELISA. Knock-down effect was determined by comparing IFNγ levels to scrambled RNA conditions. **(B)** Target cells used in A were loaded with 50uM WT-1 peptide and used as targets against WT-1 TCR T cells, knock down effect was determined by comparing IFNγ levels to that of scrambled RNA conditions. **(C)** Cell lysates MDA-MB157 and MDA-MB231, HEK293FT wt or HEK BTN3AKO, MZ1851rc and SCC9 were cultured in the presence or absence of 100 uM PAM for 24h, lysed and Western blots were probed with antibodies against candidates.

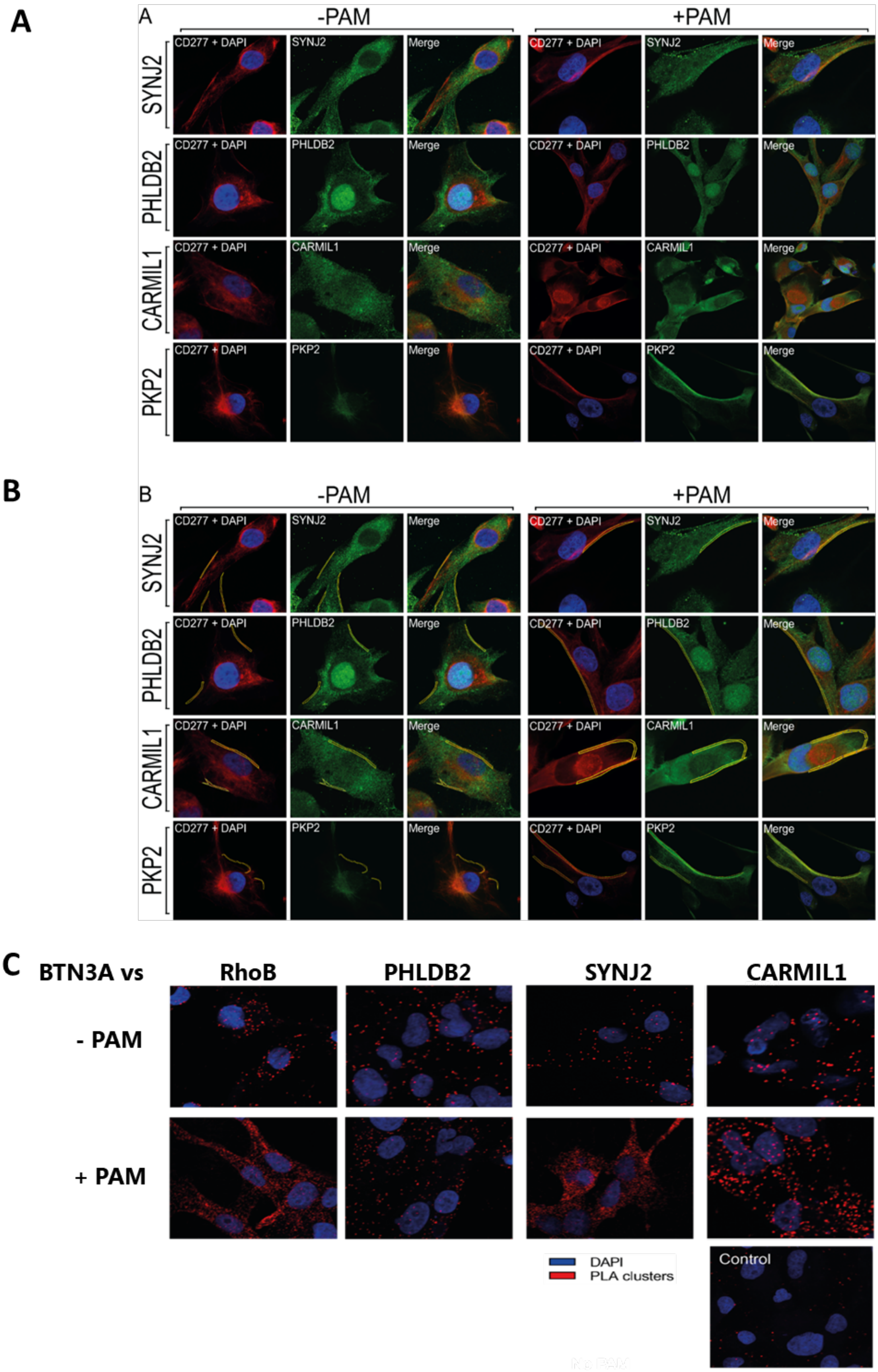


**Supplementary Figure 6. Representative images for colocalization analysis. (A)** Raw representative images for the colocalization analyses of SYNJ2, PHLDB2, CARMIL1, and PKP2 with BTN3A. **(B)** Representative images for the colocalization analyses of SYNJ2, PHLDB2, CARMIL1, and PKP2 with BTN3A with the region of interest (ROI) in yellow around the BTN3A membrane clusters. **(C)** Representative images for each of the PLA conditions are shown.
